# Reach plans for grasping vs. placing objects evoke different neural dynamics within a shared cortico-cerebellar mechanism

**DOI:** 10.64898/2026.10.14.759130

**Authors:** Petros Georgiadis, Gaelle N. Luabeya, Erez Freud, Peter J. Kohler, J. Douglas Crawford

## Abstract

Many reach studies treat grasping as the endpoint of a reach, but grasping often initiates a sequence of actions, ending with placing the object. Grasping and placing both require object-specific encoding, but placing has different computational demands, introduced by the intent, sensory feedback, and tolerance for spatial error when alignment is required. Recent neuroimaging work suggests that object placement shares similar cortico-cerebellar networks with grasp, with modest increases in network segregation and cerebellar activity. Here, we ask whether *Grasp* and *Place* are better distinguished by their spatiotemporal dynamics in both behaviour and scalp-recorded EEG activity. Using high-density EEG and motion tracking, 31 participants performed alternating *Grasp*/*Place* movements of a 3D object from/toward an oriented target/template. *Place* movements were initiated faster but executed more slowly and with greater endpoint-orientation precision than *Grasp*, indicating distinct control strategies. ERP and microstate analyses revealed that *Grasp* and *Place* shared a peri-movement polarity switch, from posterior negativity/frontal positivity during preparation to posterior positivity/frontal negativity during execution. This shared pattern was amplified during *Place* preparation and prolonged during *Place* execution, with stronger expression of preparatory medial frontal/cerebellar and execution-related posterior/cerebellar microstate dynamics. Finally, SVM decoding indicated that task-specific information was primarily encoded in low-frequency EEG activity (below 6 Hz). These results confirm that grasping and placing recruit similar cortico-cerebellar mechanisms, yet different spatiotemporal dynamics are employed to support the distinct computational demands of these behaviours.

**Significance Statement:** Grasping is often considered the endpoint of reaching, but in normal behavior, grasping is usually the first step in a sequence of object manipulations that ends with placing the object. Here, we used motion tracking and electroencephalography to directly compare spatially matched *Grasp* and *Place* movements and their cortical dynamics. We found that *Place* and Grasp utilize different control principles that are reflected in the EEG signals: a shared fronto-parietal polarity switch with differing strengths and timing. Microstate and source localization analysis further traced this to different temporal recruitment patterns of cortico-cerebellar clusters, primarily coded in low-frequency activity. These findings provide important insights into how the brain can adjust the spatiotemporal dynamics of similar networks to produce different motor behaviors.

## Introduction

Visually guided prehension requires the brain to transform object properties and location into coordinated motor commands for transporting and shaping the hand (Wolpert et al., 2001; Andersen & Buneo, 2002; Marotta et al., 2003; Castiello, 2005; Culham & Valyear, 2006; Grafton, 2010; Crawford et al., 2011). Reach-to-grasp studies have provided fundamental insight into prehension, but grasping usually supports later manipulations, often ending with placement of the object at some specific location and orientation. Our recent neuroimaging study found that placement shares considerable cortico-cerebellar topology with grasp, with subtle differences in network and cerebellar recruitment (Luabeya et al., 2026). However, it is not known how placement alters the timing of behaviour and dynamics associated with grasp, including the role of cortical oscillations.

During grasping, hand transport must be coordinated with grip formation, such that digit aperture and orientation are scaled to object features before contact (Jeannerod, 1981, 1984; Castiello, 2005). This involves both feedforward visual specification of object properties and online corrections based on sensory feedback during movement execution (Smeets & Brenner, 1999; Marotta et al., 2005; Castiello & Begliomini, 2008). Neurophysiological and imaging studies have linked these processes to a distributed visuomotor network, including posterior parietal, premotor, primary sensorimotor, and cerebellar regions (Culham & Valyear, 2006; Blohm et al., 2009; Grafton, 2010; Davare et al., 2011; Cappadocia et al., 2018; Blohm et al., 2022; Monaco et al., 2024).

Compared with grasping, object placement has received far less attention. Earlier behavioral work considered how the goal of a placing movement influences kinematic variables such as handle forces, transport speed, and movement organization (Drury & Pizatella, 1983; Quinlan & Culham, 2015; Velji-Ibrahim et al., 2022). Other studies focused on eye-hand coordination in the context of naturalistic actions with both grasp and placement (Land et al., 1999; Hayhoe, 2000; Johannsson et al., 2001; Scrafton et al., 2017). Recent behavioral work suggests that gaze has a similar influence on aiming, grasp, and placement movements (Luabeya et al., 2024), and that eye-hand coordination can be flexibly adjusted to match the demands of grasp and place (Fooken et al., 2026).

To our knowledge, only one imaging experiment has directly tested the differences between grasp and place movements in humans. In a recent functional magnetic resonance imaging (fMRI) study, we found that placement recruits similar cortico-cerebellar networks to grasp, with modest increases in cerebellar activation and functional network segregation (Luabeya et al., 2026). However, the physical and temporal limitations of this experiment did not allow measurement of detailed kinematics, neural oscillations, or real-time changes in brain activity during behavior (Tunik et al., 2008; Blohm et al., 2009; Van Der Werf et al., 2010; Verhagen et al., 2012; Schwarz et al., 2018; Guo et al., 2019; Guo et al., 2021; Xu et al., 2021; Blohm et al., 2022).

Here, we asked whether object grasp and placement recruit distinct spatiotemporal processes in the brain or modulate the timing, strength, and frequency profile of a common mechanism. To test this, we recorded hand motion and electroencephalography (EEG) signals during alternating, spatially matched grasp and place movements. To begin, we examined our kinematic data to see if *Place* alters the spatiotemporal control strategies used for grasp, specifically to optimize visual feedback. We then examined EEG signals using three methods to test whether *Grasp* and *Place* employ distinct control mechanisms or temporal modulations of a single mechanism. First, we performed event-related potential (ERP) analysis to identify task-dependent time-domain voltage differences. Second, we applied topological microstate analysis and source localization to estimate the cortico-cerebellar substrates of these differences. Finally, support vector machine (SVM) decoding across the canonical frequency bands was used to identify the spectral signature of task-related signals, and temporal generalization analysis was used to determine the temporal evolution of task-specific representations. Overall, our results supported a common spatiotemporal mechanism between grasp and placement that is differentially modulated through time for specific computational task demands.

## Materials and Methods

### Participants

Thirty-one right-handed individuals (age: 23 ± 3, 9 male) participated in the experiment. A target sample size of 30 participants was determined using a pilot-based power analysis described in the Supplementary Materials (Supplementary Figure 1), and the final EEG sample exceeded this target by one participant. ERP, microstate, and SVM decoding analyses were conducted on the complete EEG dataset after preprocessing and participant-level quality control (N = 31). Of these 31 participants, kinematic analyses included 25 with complete, usable motion-tracking data; we excluded six because of incomplete motion-capture recordings or excessive marker loss. Electrooculography (EOG)-based eye-movement onset and eye-hand timing analyses included 24 participants after excluding participants with insufficient valid EOG trials or unreliable onset estimates due to noisy periocular channel recordings. Handedness was self-reported and confirmed using the Edinburgh Handedness Inventory (EHI) (all scored above 52, mean: 85.45 ± 15.3; Oldfield, 1971). All participants reported normal or corrected-to-normal vision and no known neurological disorders on the day of the experiment. Participants were recruited from pools of York University trainees and provided written consent prior to their participation. All experimental procedures were approved by the York University Human Participants Review Committee (HPRC: 2023-117).

### Experimental Setup

The experimental setup is shown in Figure 1A. Experiments were performed within an isolated room, custom-insulated for EEG recordings. Participants were seated comfortably with their heads stabilized using a chin rest. The right hand initially rested on a “home” button positioned on a table, immediately in front of their chest. The grasp/placement object (a 10 x 3 x 2 cm rectangular plastic block) was initially placed within a template mounted on an inclined board, approximately 40 cm in front of the participant, within comfortable reaching distance. The template was designed to loosely but precisely hold the block, without interfering with manual interaction. The template in turn was mounted on a slot that allowed changes in location (± 15cm) and orientation (± 30°) against stoppers at either end of play. The trajectory of the slot was angled to match the biomechanics of lower arm rotation. An experimenter behind the board controlled the location and orientation of the template as described below. Three LEDs were mounted on the tips of adjustable rubber/fibreglass ‘snakes’ above the board: two white LEDs were positioned laterally to illuminate the object/template at its left and right positions, while a central red LED served as the gaze fixation point for the start of each trial.

**Figure 1.**
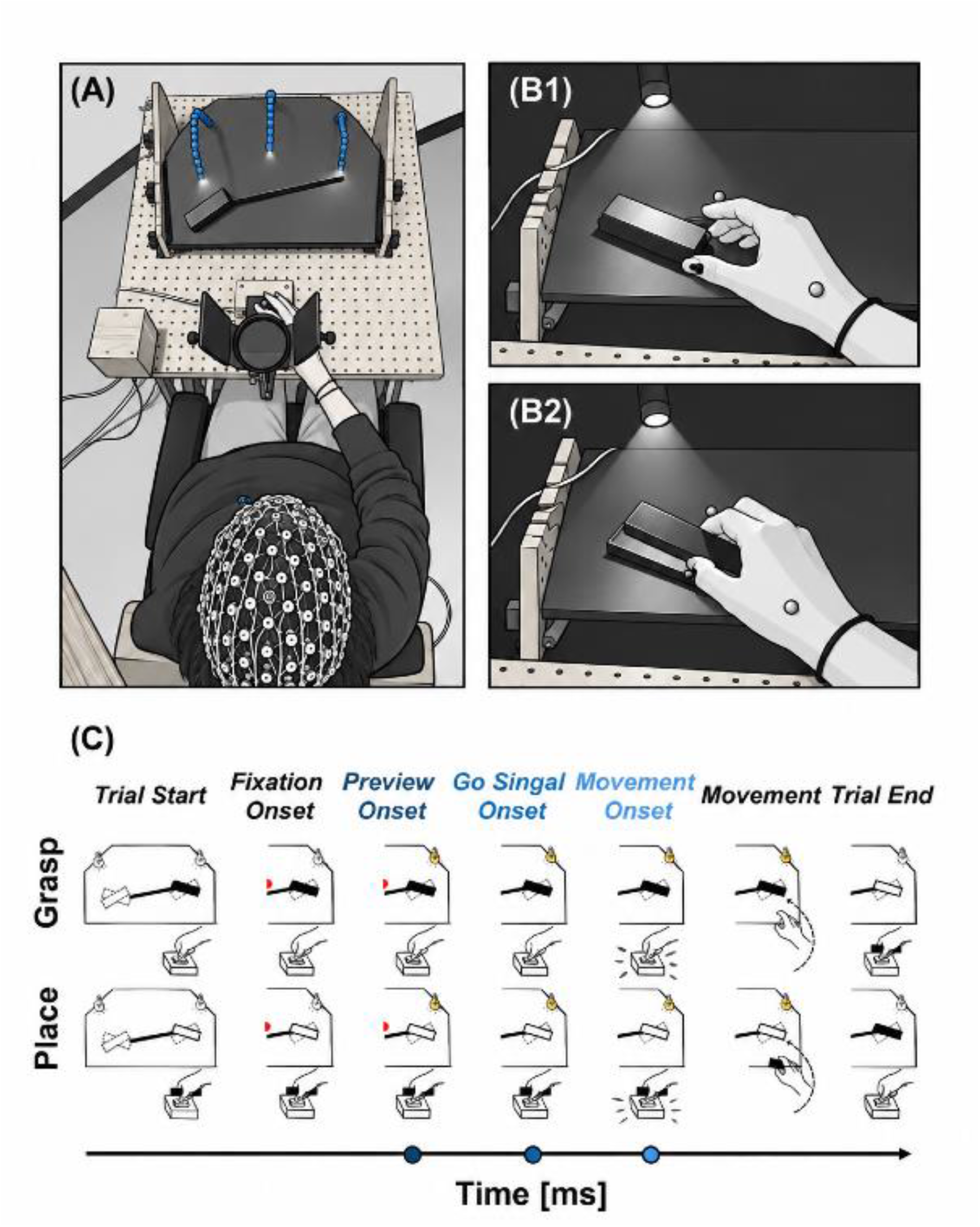
Experimental setup for the *Grasp* or *Place* task. **A**. A bird’s-eye view of the apparatus set up in the EEG chamber. The participant is comfortably seated on the chair with their hand on the “home” button. Immediately in front of them is the chinrest, where they kept their heads fixed during the experiment, and the apparatus with the blue rubber/fiberglass “snakes” hosting the LED lights (left, right illuminators, and a central fixation light) and the inclined board with the object and template. **B**. A representation of the participant’s point of view during movement execution for the 2 main phases of the experiment, *Grasp* (**B1**) and *Place* (**B2**). During the experiment, the only light inside the EEG chamber came from these three illuminators. **C**. The progression of the experimental *Grasp* and *Place* trials. Each trial began with the participant pressing and holding a “home” button in complete darkness. A central red LED was then illuminated to enforce fixation. After 1 s, an illuminator above the object location turned on, providing peripheral information about object location and orientation while fixation was maintained for a variable delay (1000-1500ms). The extinction of the central LED served as the “GO-signal”, upon which participants initiated their movements about 450ms after (mean across conditions), while the object illuminator remained on to allow for visually guided execution of the movement. Movement onset was recorded upon “home”-button release. In *Grasp* trials, participants grasped the object and returned to the “home” position while holding it. In the following *Place* trial, participants placed the object into the template and returned to the “home” position without the object, preparing for the upcoming *Grasp* trial.

### Experimental Design

Participants completed two experimental sessions on separate days. Sessions were scheduled at approximately the same time of day, with no more than one week between sessions. Each session consisted of eight blocks, with 64 trials in each block, for a total of 512 trials per session. Within each block, *Grasp* trials and *Place* trials alternated, such that a *Grasp* trial (a reach from “home” position to acquire the block) was always followed by a *Place* trial (a reach from “home” position to replace the block into the template). Two object locations (15cm left, right) and two object orientations (30° clockwise, 30° counterclockwise) were alternated, following a randomized schedule, to maintain task attention and reduce potential adaptation effects in neural responses. This resulted in a 2 (location) × 2 (orientation) × 2 (task) within-subject design. Within each block, the object/template’s location and orientation were randomized. One experimenter monitored the EEG recording from within the EEG chamber throughout the session, and another manually adjusted the object template position and orientation (by sliding & rotating the template against stoppers) before each trial. Instructions regarding the required configuration were delivered via headphones and were audible only to the experimenter. Experiments were performed in complete darkness except for the stimulus events described below.

The trial sequence is illustrated in Figure 1C. Each *Grasp* trial (upper row) began with the participant pressing and holding the “home” button. A central red LED was then illuminated, which participants were instructed to visually fixate. After 1 s, the LED illuminator above the object location was turned on, providing visual information about the object’s location and orientation, while fixation on the red LED was maintained. This preview period lasted for a variable delay of 1000-1500 ms to mitigate anticipatory movements. At the end of this delay, the central red LED was extinguished, serving as the “GO”-signal for the participant to initiate their movement. The illuminator above the object location remained on, allowing participants to visually guide their movement. Participants initiated the movement by releasing the “home” button and were allowed to move their gaze freely. After grasping the object (typically with a pinch grip, using their thumb and index fingers), they returned to the “home” position while maintaining the object in their hand. With the re-pressing of the “home” button, the illuminator turned off, and the *Grasp* trial was complete.

After readjustment of the template by the experimenter (to the same or different location/orientation), the subsequent *Place* trial began, following a similar sequence and the same timing (Figure 1C, bottom row). Note that this time the trial starts with participants pressing the “home” button with the object in their hand. In this case, participants had to orient the object precisely to fit it into the template. After placing the object in the template, they then returned the hand to the “home” position (without the object) and pressed the button again to prepare for the next *Grasp* trial. In this way, we were able to record a natural sequence of reach-grasp-place movements and obtain the maximal amount of spatially matched *Grasp* and *Place* trials during the recording period.

### Data Acquisition

ELECTROENCEPHALOGRAPHY. Data acquisition was controlled by a second experimenter located in a control room adjacent to the experiment room. EEG was recorded using a 128-channel HydroCel Geodesic Sensor Net (GSN; HydroCel GSN 129, Electrical Geodesics Inc.). During acquisition, EEG data were referenced to the vertex electrode located on the midline over the motor cortex, as implemented in NetStation software (version 5.5) for EEG acquisition and recording (Electrical Geodesics, Inc.). Electrode impedances were maintained below 5 kΩ throughout recording. Signals were sampled at 1000 Hz and band-pass filtered between 1 and 50 Hz to attenuate slow drifts and high-frequency noise. Line noise was further reduced using a 60 Hz notch filter.

BEHAVIOURAL DATA. Kinematic data were recorded simultaneously using an OptiTrack motion capture system (NaturalPoint Inc., USA) at 120 Hz through the Motive software (version 3.3.0; NaturalPoint Inc.) for motion tracking and data acquisition. Reflective markers were placed on the dorsal surface of the distal phalanges of the right thumb and index finger, as well as on a midpoint between the trapezium and trapezoid bones, on the dorsal area of the hand. This configuration defined a three-point plane, allowing reconstruction of the midpoint and finger positions relative to the reference point placed at the right corner of the experimental table, proximal to the participant. Using the location of the midpoint allowed for estimating the position and velocity of the hand, while using the locations of the 2 fingers allowed for estimating the grip aperture and orientation of the precision grip. EOG activity was derived from six periocular EEG electrodes: E128 and E125 positioned lateral to the left and right eyes, respectively; E21 and E14 positioned above the eyes; and E127 and E126 positioned below the left and right eyes, respectively. These signals were used to estimate the onset and duration of eye movements.

SYSTEMS SYNCHRONIZATION. A custom MATLAB script (MathWorks, Natick, MA) controlled a National Instruments (NI) interface to coordinate communication between NetStation (EEG acquisition) and Motive (OptiTrack acquisition) interfaces. This setup ensured precise synchronization between EEG / EOG and kinematic recordings and enabled the logging of events. Motion capture recording was initiated at trial onset, defined as the moment the participant pressed and held the “home” button; this event was simultaneously registered in NetStation. Illumination of the object location was logged as a separate preview-onset event. The extinction of the central red LED was also logged in NetStation as the “GO”-signal. Movement onset was defined by the “rising edge” signal, associated with the release of the “home” button. Trial offset was recorded when participants returned to and pressed the “home” button after completing the movement.

### Data Preprocessing

KINEMATICS. Kinematic data were preprocessed using custom MATLAB scripts. Trial-wise kinematic files were matched to experimental conditions, and missing samples were interpolated using shape-preserving cubic interpolation. Marker trajectories were then low-pass filtered (4th-order zero-phase Butterworth, 6 Hz cutoff) to reduce high-frequency noise (Winter, 2009). Movement onset was identified as the first sample at which resultant hand speed across the three axes exceeded 5% of the trial-specific peak speed; movement offset was identified as the first post-peak sample at which speed fell below the same threshold, corresponding to object contact in *Grasp* trials and template contact in *Place* trials. Reaction time (RT) (time from “GO”-signal to “home”-button release) and movement duration (MD) (time from movement onset to completion touch) were calculated. Trials with reaction times smaller than 100 ms or exceeding 1000 ms were discarded from all further analysis (mean = 4.61%, SD = ± 4.10%). Peak velocity (PV) was calculated as the maximum speed of the hand’s marker between movement onset and contact. Endpoint pinch-grip orientation was quantified at this time point using the vector from the thumb marker to the index-finger marker. This vector was converted to spherical coordinates to obtain azimuth and elevation angles, with the thumb-index distance retained as the vector magnitude (grip-aperture). Grip-orientation angles were then expressed relative to the instructed object/template orientation, such that values reflected normalized endpoint-orientation deviations from the object’s/template’s orientation.

EEG SIGNALS. EEG data were preprocessed in MATLAB using EEGLAB (Delorme and Makeig, 2004). Continuous EEG recordings were imported, downsampled to 100 Hz, band-pass filtered (1-50Hz), and cleaned for line noise. Non-EEG channels (from the face and neck area) were excluded from the main EEG dataset and retained separately when needed for artifact identification. Prior to independent component analysis, the continuous EEG data were denoised using the Generalized Eigenvalue De-Artifacting Instrument (GEDAI) approach, which uses generalized eigenvalue decomposition to separate activity consistent with plausible brain-generated EEG from spatially implausible artifact-dominated activity (Ros et al., 2025). GEDAI was used conservatively as an initial spatial denoising step to reduce large-amplitude artifacts before ICA, and the resulting data were visually inspected to ensure preservation of physiologically plausible EEG signals.

Bad (noisy or unresponsive) channels were identified using probability, kurtosis, and spectral criteria, supplemented by visual inspection, and were temporarily removed before ICA. The rank of the cleaned EEG data was estimated, and ICA was performed with rank-adjusted dimensionality reduction where necessary. Components reflecting ocular, muscle, or line-noise artifacts were identified using automated IC classification (ICLabel; Pion-Tonachini et al., 2019) and visual inspection of their scalp maps, time courses, and spectra, and were removed from the data. Removed channels were subsequently interpolated using a three-dimensional spline procedure (Perrin et al., 1987), and the data were re-referenced to the average reference. Cleaned continuous data were then epoched around movement onset and imported to (a) LIMO M/EEG toolbox (Pernet et al., 2011) for inspecting task-effects at the ERP-level and (b) Cartool for testing microstate-based segmentation and template-map backfitting analyses. Figure 2 depicts the EEG analysis pipeline after the preprocessing was applied.

**Figure 2:**
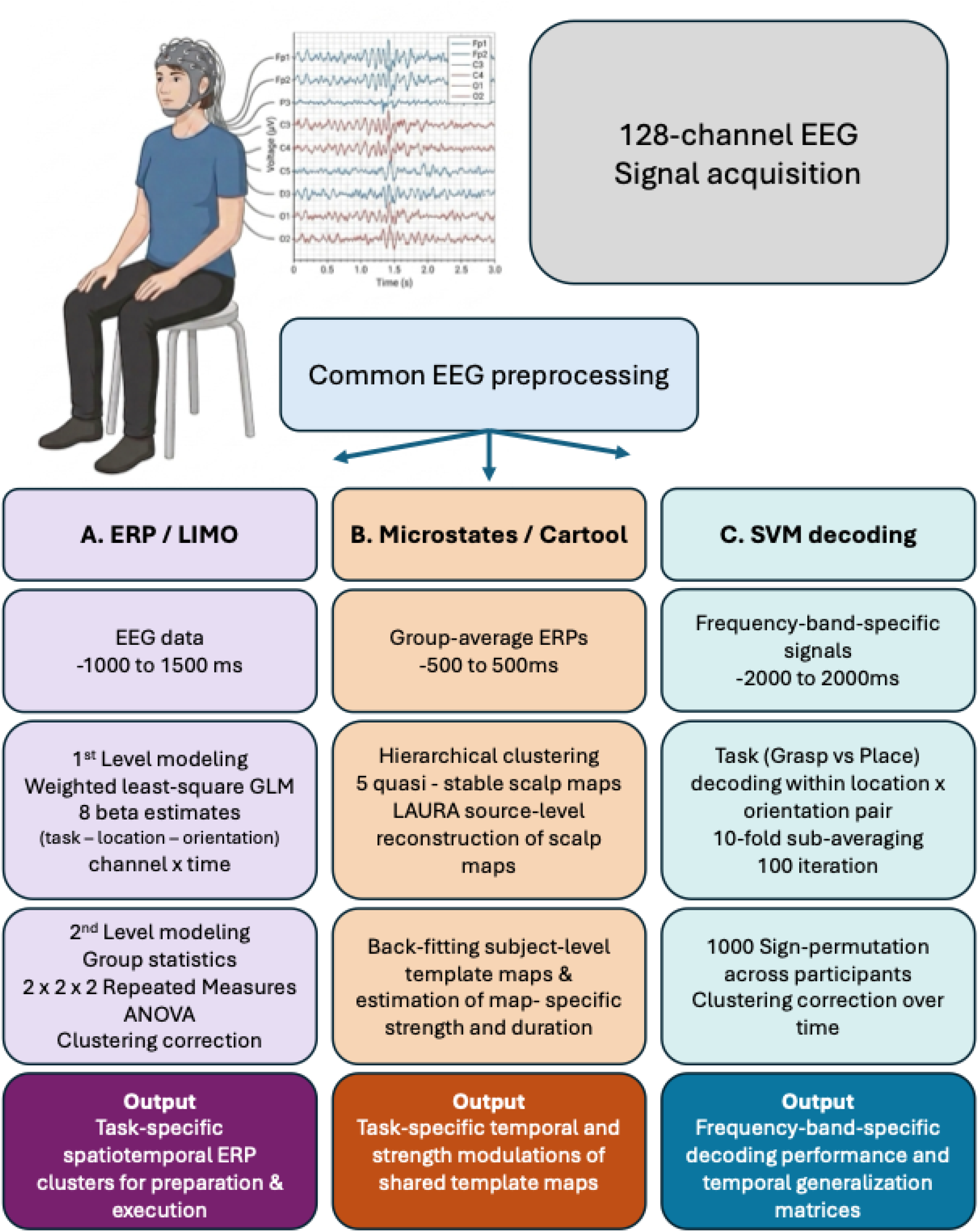
Schematic overview of the EEG analysis workflow used to compare *Grasp* and *Place*. Following 128-channel EEG signal acquisition, data underwent a common preprocessing pipeline (see Data “Preprocessing/EEG Signals”) before being submitted to three complementary analysis branches. **A**, The ERP/LIMO branch used movement-onset-locked EEG epochs from −1000 to +1500 ms, first-level weighted least-squares general linear models, and group-level 2 × 2 × 2 repeated-measures ANOVAs with cluster correction to identify task-specific spatiotemporal ERP clusters. **B**, The microstate/Cartool branch used group-average ERPs from −500 to +500 ms to identify five quasi-stable scalp maps, estimate their source-level organization with LAURA, and quantify subject-level map-specific strength and duration through back-fitting. **C**, The SVM decoding branch used broadband and frequency-band-specific EEG signals from −2000 to +2000 ms to classify *Grasp* versus *Place* within matched location × orientation pairs across time, followed by participant-level sign-permutation testing and cluster correction over time, to identify temporal clusters of significant above-chance (50%) performance. Together, these analysis branches tested task differences in ERP amplitude, scalp-map dynamics, and frequency-specific multivariate decoding performance.

EOG. To assess whether task-related EEG effects were accompanied by systematic differences in eye movements induced by the task (Voudouris et al., 2018; Fooken et al., 2026), we analyzed EOG activity in a separate preprocessing branch. EOG provides an indirect measure of eye movements by recording voltage changes generated by the corneo-retinal standing potential, with horizontal and vertical eye movements typically captured using lateral canthal and supra-/infra-orbital electrode pairs, respectively (Creel, 2019; Chang et al., 2019). Periocular electrodes were therefore retained from the HydroCel net (E128, E125, E21, E14, E127, and E126) and used to derive bipolar vertical and horizontal EOG signals. Data were filtered from 0.1 to 15 Hz, cleaned for line noise when applicable, and downsampled to 100 Hz using EEGLAB (Delorme and Makeig, 2004). Vertical EOG was calculated separately for the left and right eyes as upper-minus-lower periocular activity, and the two signals were averaged to obtain a single VEOG trace. Horizontal EOG was calculated as the difference between left and right lateral periocular electrodes; because the direction of horizontal gaze shifts was not of primary interest, the absolute HEOG signal was used for task comparisons. Individual trials were rejected from both EOG and EEG further analyses if the mean amplitude of any retained EOG/frontal channel exceeded ±100 μV in a preview-period inspection window. The remaining trials were baseline-corrected using a trial-specific baseline during central fixation for both the vertical and horizontal components of EOG. Task effects in VEOG and absolute HEOG were later assessed over the peri-movement interval using time-resolved paired comparisons with cluster correction across time.

### Statistical Analysis

KINEMATICS. For the kinematic analysis, participant-level means were first computed separately for *Grasp* and *Place* trials for each dependent measure: RT, MD, PV, and endpoint grip-orientation variability. Group-level task effects were tested using paired-samples t-tests between *Grasp* and *Place*-specific measures across participants. Prior to statistical testing, the assumption of normality was confirmed using the Shapiro-Wilk test (Shapiro & Wilk, 1965). Effect size was quantified using Cohen’s d. The α-level for statistical significance was set to 0.05. All statistical analyses were performed in R (R Core Team, 2025).

EOG. To evaluate whether eye movements influenced our EEG data, we aligned EOG signals to the hand-movement onset; i.e., the same event alignment used for all EEG data analysis. For this, trial-level VEOG and HEOG signals were analyzed within a peri-movement window from −500 to +500 ms relative to movement onset (similar to the one used for “Microstates” analysis). VEOG was analyzed as a signed signal to preserve the direction of vertical eye movements. HEOG was transformed to absolute amplitude before statistical testing, because leftward and rightward eye movements can produce opposite polarities; this analysis therefore tested task differences in HEOG magnitude rather than direction. For each participant, task, signal, and time point, mean EOG amplitude was first computed across trials. Subject-level task contrasts were then calculated at each time point using effect coding for Task, with *Grasp* coded as −1 and *Place* coded as +1, such that positive contrast values indicated greater EOG amplitude for *Place* than *Grasp*. Group-level task effects were assessed separately for VEOG and absolute HEOG using one-sample t-tests against zero across participant-level task contrasts at each time point.

To correct for multiple comparisons over time, statistical significance was assessed using a cluster-based sign-flip permutation procedure. Contiguous time points exceeding an uncorrected cluster-forming threshold of p < 0.05 were grouped into temporal clusters, and cluster mass was defined as the sum of the absolute t values within each cluster. The null distribution of maximum cluster mass was estimated using 5000 random sign-flip permutations across participants. Clusters were considered significant if their permutation-corrected p-value was below α = 0.05. Supplementary Figure 2C demonstrates the EOG group-level activity within the −500 to +500 ms of analysis, relative to movement onset, along with the task-differentiating temporal clusters. For HEOG, the significant cluster extended from 170 to 500 ms after hand movement onset (p = 0.014). For VEOG, the significant cluster extended from 190 to 500 ms after hand movement onset (p < 0.001). No significant task differences were observed before movement onset. This suggests that the gaze data reported below in Results did not affect our main reach task analysis.

LIMO M/EEG ERP ANALYSIS. Task-related ERP activity was analyzed using the LIMO M/EEG toolbox (Pernet et al., 2011). For each participant, movement-onset-locked EEG epochs from −1 to +1.5 s were entered into a mass-univariate general linear model (Figure 2A). First-level models were fitted independently at each retained electrode and time point, estimating beta coefficients for each of the eight experimental conditions (task by location by orientation). Model parameters were estimated using weighted-least-squares (WLS), which reduces the influence of outlier trials by down-weighting noisy observations (Pernet et al., 2022). This produced subject-specific beta estimates for each condition, channel, and time point.

Group-level effects were tested using repeated-measures ANOVAs with the within-participant factors of task, location, and orientation. Separate 2 × 2 × 2 ANOVAs were computed at each retained electrode and time point, producing mass-univariate statistical parametric maps for each main effect and interaction. Statistical inference used robust estimates based on 20% winsorized means and a generalized Welch’s *F* statistic, reducing sensitivity to unequal variances and outliers (Delacre et al., 2019; Pernet et al., 2011). Multiple comparisons across electrodes and time points were controlled using non-parametric bootstrap spatiotemporal clustering with 1000 iterations and a family-wise α level of 0.05 (Maris and Oostenveld, 2007). Cluster-corrected results are reported for the main effect of task, contrasting *Grasp* and *Place* movements.

MICROSTATE ANALYSIS. To examine whether *Grasp* and *Place* movements differed in the sequence and expression of large-scale scalp potential configurations, we conducted a topological microstate analysis on movement-onset-locked ERPs using Cartool (version 3.33; Denis Brunet; http://brainmapping.unige.ch/Cartool.htm) (Figure 2B). This approach evaluates changes in the spatial organization of the scalp electric field rather than focusing on voltage changes at individual electrodes (Murray et al., 2008). Because scalp topologies are constrained by the configuration of the underlying neural generators, differences in the timing or prevalence of specific maps are treated as differences in the functional neural states engaged by each task. Microstate analysis has previously been used to show that visually guided grasping and online movement adjustments involve differential temporal recruitment of distinct parietal brain states (Tunik et al., 2008).

For the very first step of the microstate analysis, we computed the group-averaged ERPs from the experimental conditions within a peri-movement window from −500 to +500 ms relative to movement onset. These were then subjected to the hierarchical clustering procedure (Brunet et al., 2011), implemented in Cartool, to identify the dominant scalp maps that summarized the main topologic patterns present across conditions and time. The number of maps retained for further analysis was selected using Cartool’s optimization criteria, including the modified Krzanowski-Lai criterion and the global explained variance. This procedure identified five template maps that captured the dominant topologic structure of the ERP data during the transition from movement preparation to execution.

These group-level template maps were then fitted back to the ERP data of each participant and condition. For every time point, the participant-level scalp field was assigned to the template map with the highest spatial similarity. This back-fitting step provided subject-level estimates of how often, how strongly, and how well each map characterized the ERP activity in each task condition.

For each participant, task, and template map, we extracted the measures of time coverage and mean global field power. Time coverage (%) measured the proportion of the analyzed epoch assigned to a given map for each subject-specific ERP. Mean global field power (GFP; i.e., the spatial standard deviation across electrodes) quantified the strength of the scalp field during the periods assigned to a given map at the subject-level ERPs. These measures were analyzed using Repeated-Measures ANOVAs to test whether *Grasp* and *Place* differed in the recruitment of specific microstate configurations.

To aid anatomical interpretation, the template maps were also submitted to distributed source estimation using the Local Auto-Regressive Average (LAURA; Menendez et al., 2001) inverse solution in Cartool. Source estimates were computed in standard MNI anatomical space and summarized using the AAL3 atlas (Rolls et al., 2020). For each map, regional maxima and ranked source contributions were extracted descriptively to identify the cortical regions most strongly associated with the observed scalp configurations.

SUPPORT VECTOR MACHINE DECODING. Following the ERP and microstate analyses, time-resolved multivariate decoding was used to identify the spectral range carrying task-specific information in the EEG signal. Decoding was performed separately for each participant using linear support vector machines (Chang and Lin, 2011) implemented in scikit-learn and MNE-Python. The same movement-onset-locked EEG trials from −2 to +2 s were analyzed here at 200 Hz. Activity was represented in non-overlapping 10-ms spatiotemporal windows (similar to Guo et al. (2019)), yielding 198 features per window. Feature standardization was estimated from the training data only.

To prevent the levels of the Location (left, right) and Orientation (CW, CCW) factors from contributing to Task classification, *Grasp* and *Place* trial numbers were matched within each Location × Orientation condition and divided into non-overlapping pseudotrials. Cross-validation left one pseudotrial from each of the eight Task × Location × Orientation conditions out for testing on each fold, thereby maintaining identical Location × Orientation composition between *Grasp* and *Place* in both training and testing sets. To characterize the spectral profile of task-discriminative information, we followed the frequency-specific decoding procedure demonstrated by Bednar and Lalor (2018). Decoding performance was repeatedly estimated across overlapping 2-Hz frequency ranges from 1-30 Hz, advanced in 1-Hz steps, while all other decoding parameters were held constant. Frequency-specific decoding was evaluated across time, with mean accuracy used to summarize each frequency range. Group-level decoding was assessed against 50% chance using two-tailed cluster-based sign-permutation tests across participants (2,000 permutations), with a cluster-forming threshold of p < 0.05 and cluster-corrected significance at p < 0.05.

TEMPORAL GENERALIZATION. To characterize the temporal stability of task-discriminative EEG patterns, we performed temporal generalization analysis (King & Dehaene, 2014; Guo et al., 2019). Using the same pseudotrial construction, feature standardization, cross-validation scheme, and linear SVM described above, a classifier was trained independently at each 10-ms window and tested at every other 10-ms window across the −2 to +2 s movement-onset-locked epoch. This yielded a 400 × 400 training-time × testing-time decoding matrix for each participant, with classification accuracy expressed relative to 50% chance. Group-level inference was performed across the training × testing matrix using two-tailed cluster-based sign-permutation tests across participants (2,000 permutations). Clusters were formed from neighboring train-test samples exceeding a two-sided cluster-forming threshold of p < 0.05 and were considered significant at cluster-corrected p < 0.05.

## Results

The goal of this study was to determine if placement and grasping are implemented through distinct control mechanisms, or through spatiotemporal modulations of an otherwise common mechanism. We tested this question at four levels. First, we analyzed behavioral kinematics to determine if our two tasks employ different spatiotemporal control strategies. Second, we employed ERP analysis to test whether scalp-level voltage fluctuations differ spatially or temporally between the two tasks. Third, we used microstate analysis along with source localization to estimate the cortico-cerebellar substrates of these task-specific signals. Finally, we used SVM decoding and temporal generalization to determine the spectral profile of the task-discriminatory signals and temporal stability of the task-specific representations across movement preparation and execution.

### Behavioral Kinematics

First, we examined our kinematic data to see if it would provide clues to the control strategies used in our two tasks. Specifically, we tested the hypothesis that our *Place* task required increased online control to match the object and template, whereas small variations in hand orientation and finger configuration could still provide a successful *Grasp*. This hypothesis leads to the prediction that the *Place* task would lead to slower but more precise movements than *Grasp*.

TEMPORAL ANALYSIS. Figure 3A contrasts the velocity profiles of the moving hand across our two tasks, from movement onset to contact. This visualization already indicates clear task-specific differences in the temporal progression of visually guided *Grasp* and *Place* tasks. *S*pecifically, participants initiated reach movements significantly faster during *Place (M = 0.436 s)* compared to *Grasp (M = 0.460 s) (t(24) = 3.03, p = 0.0058, Cohen’s d = 0.61),* but movement duration was significantly longer for *Place (M = 1.122 s*) than *Grasp* movements *(M = 0.815 s) (t(24) = −9.38, p = 1.70 × 10⁻⁹, Cohen’s d = −1.88).* Likewise, Peak velocity was significantly lower for *Place (M = 0.451)* compared to *Grasp (M = 0.526) (t(24) = 5.50, p = 1.18 × 10⁻⁵, Cohen’s d = 1.10).* In summary, *Place* movements started sooner, were slower, and lasted longer.

**Figure 3:**
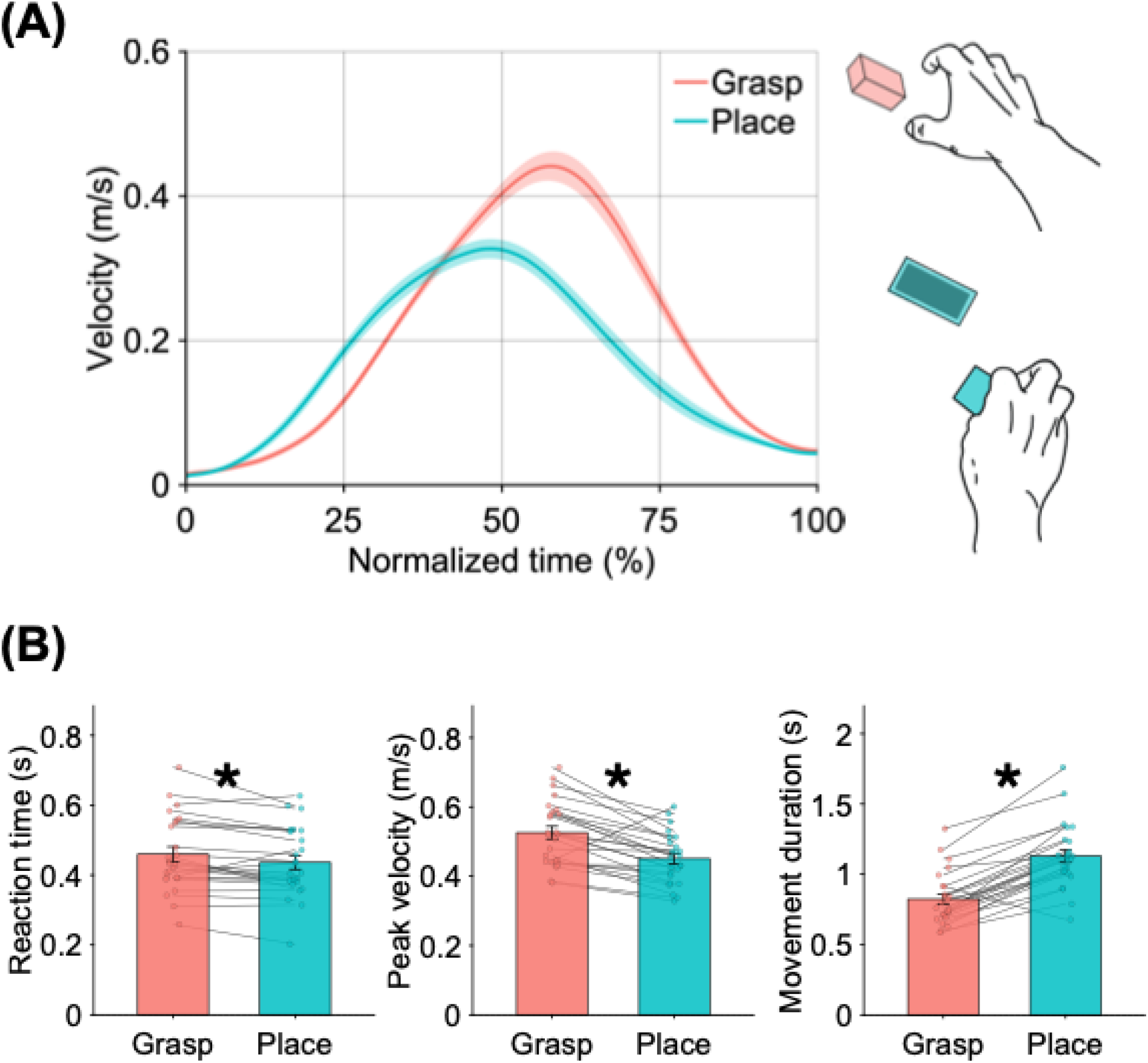
Temporal analysis of hand movements. **A.** Group-average velocity profiles for *Grasp* and *Place* trials. Velocity was computed from the hand marker trajectory and time-normalized from movement onset to object contact, with the x-axis indicating normalized movement time (%). Shaded regions indicate SEM across participants. *Grasp* movements showed a later and higher velocity peak, whereas *Place* movements showed a broader, lower-amplitude profile. **B.** Participant-level kinematic measures for reaction time, peak velocity, and movement duration. Bars show group means, error bars indicate SEM, dots show individual participant means, and gray lines connect paired *Grasp* and *Place* values from the same participant. *Place* movements had shorter reaction times, lower peak velocity, and longer movement duration than *Grasp* movements. Asterisks indicate significant paired-samples t tests between tasks.

Eye movements preceded the hand movement but otherwise followed a similar time course as the *Grasp* and *Place* hand movements (Supplementary Figure 2). To quantify gaze timing, eye-movement onset was estimated on each trial as the first time point at which the combined EOG amplitude, defined as √*VEOG^2^* + *HEOG^2^*, exceeded 5% of the trial-specific peak EOG amplitude. Eye-movement onset differed significantly between tasks, occurring later for *Place* than *Grasp* trials relative to the “GO” Signal (*Grasp* - *Place*: −9.96 ms; paired t-test: t(23) = −2.77, p = 0.01). The relative timing between eye and hand movement onset also differed significantly between tasks (*Grasp* - *Place*: −23.7 ms; t(23) = −3.46, p = 0.002). Thus, although *Place* hand movements started earlier than *Grasp* movements, the eye movements associated with *Place* were initiated later relative to the hand, indicating a change in eye-hand coordination.

SPATIAL ANALYSIS: REACH ERRORS. A significant task effect was also observed for our precision measure. Figure 4 demonstrates the orientation of the end-point pinch grid for the 2 tasks. Endpoint grip-orientation variability was significantly lower for *Place (M = 33.44°)* compared to *Grasp (M = 37.03°) (t(24) = 2.41, p = 0.024, Cohen’s dz = 0.48)*, indicating that *Place* movements produced more consistent end-point grip orientations than *Grasp* ones. In contrast, the magnitude of the absolute orientation error relative to the target orientation did not differ significantly between the tasks. Thus, *Place* improved endpoint orientation precision, but not orientation accuracy. The Euclidean end-point location error was also investigated for task-specific differences, but the 2 tasks did not differ with regard to the magnitude or the variability of this error measure (Supplementary Figure 3).

**Figure 4:**
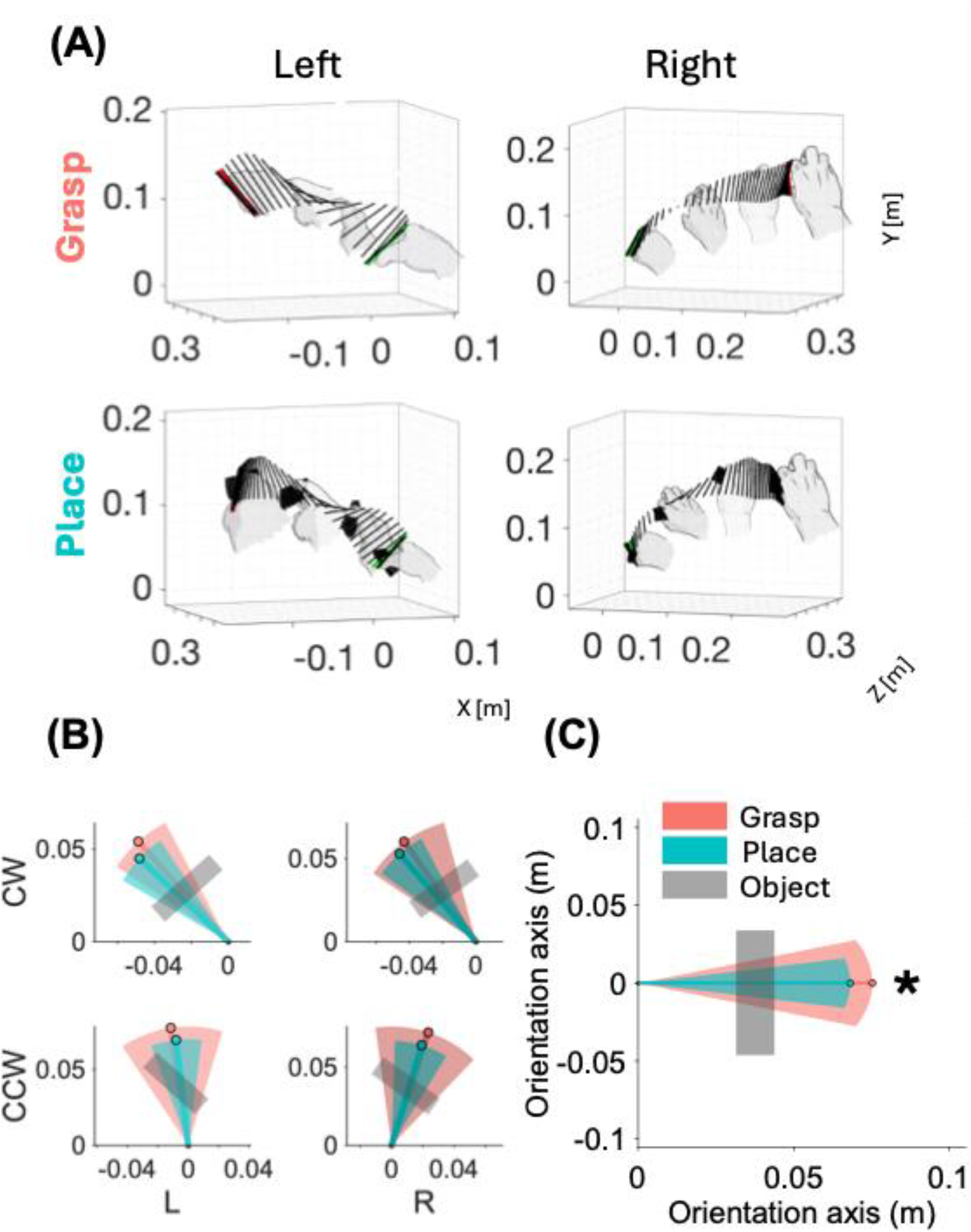
Spatial analysis of hand movements. **A**, Kinematic trajectories for a representative trial of each task-location-orientation condition. *Grasp* trials are shown in the upper row; *Place* trials in the lower row. Leftward movements are displayed in the left column, rightward movements in the right column, all for a counterclockwise orientation. The 3D plots (x: lateral displacement, y: height, z: depth) depict the trajectory of the pinch grip through space and time, with lines connecting the thumb and index finger plotted every 10 frames. The first time-point is highlighted in bold green, and the final time-point (time of object contact or placement) in bold red. Hand illustrations are overlaid to aid interpretation of the movement. **B**, Endpoint pinch-grip orientation shown separately for left and right target locations and for clockwise and counterclockwise object/template orientations. Lines show the group-mean thumb-to-index vector for each task, with line length reflecting mean grip aperture. Shaded wedges indicate task-level angular variability, computed by rotating the mean grip vector by ±1 circular standard deviation. Thick semi-transparent gray bars indicate the instructed object/template orientation. *Grasp* and *Place* are colour-coded. **C**, Task-level endpoint-orientation variability after condition-centering. Subject-level grip orientations were first realigned relative to the corresponding task, location, and object-orientation mean, and then pooled across location and orientation conditions. Lines show the task-level mean vector, line length reflects mean grip aperture, and shaded wedges show task-level angular variability. The thick semi-transparent gray bar indicates the reference object/template orientation. *Place* movements showed reduced endpoint-orientation variability relative to *Grasp*. Asterisks (*) indicate significant paired-samples t tests.

### EEG

We next tested whether the behavioral differences we identified between *Grasp* and *Place* were associated with distinct neural configurations in our EEG dataset or changes in the timing and strength of shared peri-movement dynamics. Once again, this was a three-stage approach: first on spatiotemporal contrasts at the scalp level, then on localizing the source of these patterns, and then on frequency-specific oscillations.

SPATIOTEMPORAL ANALYSIS: EVENT-RELATED POTENTIALS. We examined scalp-level ERPs to see if our tasks evoked different spatial or temporal patterns. Based on our previous fMRI results (Luabeya et al. 2026), we expected similar spatial patterns. Based on our kinematic results, one might reasonably expect the timing of placement signals to be extended (to match the behavior) and the relative power of these patterns to be modulated to match the increased visual and precision demands of the place task.

To get a sense of the ERPs evoked by the two tasks, we first inspected the activity patterns of electrodes in the contralateral hemisphere, grouped into families of frontal, central, parietal, and occipital electrodes (see Figure 5A). We then computed task-specific grand-average ERPs for each electrode and plotted them grouped by family in Figure 5B, while Figure 5C presents the whole-scalp topologies of the ERPs at five key time points spanning from late movement preparation to early movement execution. This was done for illustration purposes and did not serve as input to subsequent analyses. The two tasks showed highly similar scalp-level ERP dynamics, with differences mainly reflecting the strength and duration of shared voltage patterns rather than distinct topological configurations. Both showed a ‘switching’ pattern from negative (blue) occipital-parietal / positive (red) central-frontal signals during planning (before movement onset) to positive occipital-parietal / negative central-frontal during early execution, as expected from previous studies (Xu et al., 2021; Pereira et al., 2021).

**Figure 5:**
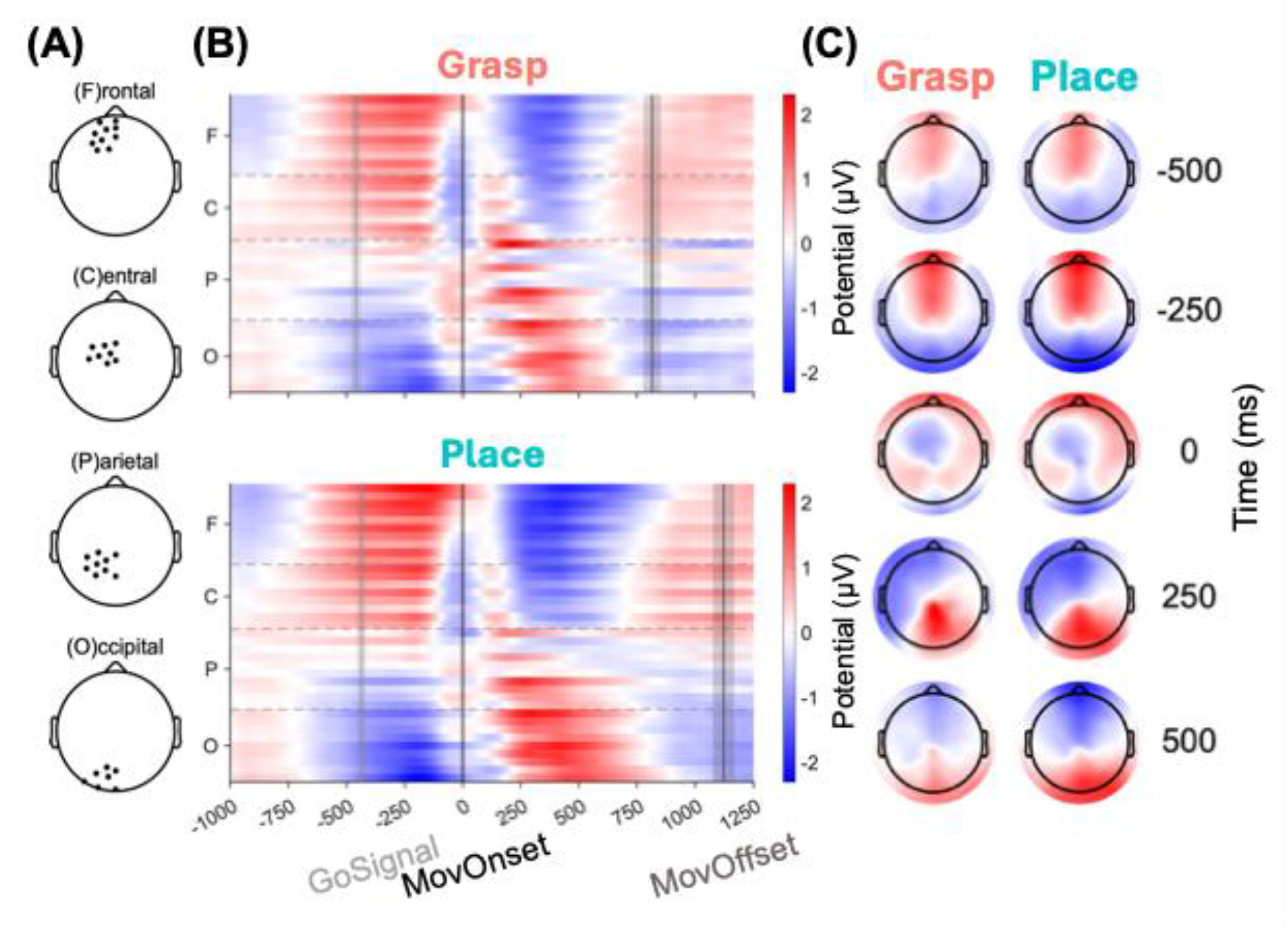
Grand-average ERPs for *Grasp* and *Place* trials, shown for contralateral frontal, central, parietal, and occipital electrode families (y-axis, top to bottom). **A,** Topological representation of the channels included in each channel family: (F)rontal, (C)entral, (P)arietal, and (O)ccipital, projected on the y-axis of the ERP plots. **B**, Grand-average ERPs for *Grasp* (upper panel) and *Place* (lower panel). Each horizontal band represents the ERP activity from one electrode in the respective family, with dashed horizontal lines separating electrode families. ERPs are time-locked to movement onset, with vertical markers indicating the “GO”-signal (light-gray), movement onset (black), and movement offset (dark-gray). **C**, Scalp topologies show the corresponding voltage distributions at five representative time points, time-locked to movement onset (−500, −250, 0, +250, and +500 ms), for *Grasp* (left) and *Place* (right). Colour scales indicate ERP amplitude in microvolts, with red (+) and blue (−) denoting opposite voltage polarities.

To quantify subtle task-related differences in the ERPs, we computed first-level condition-specific beta estimates for each participant (see Methods section *Statistical Analysis*; LIMO M/EEG ERP ANALYSIS), and analyzed them at the group level using mass-univariate repeated-measures ANOVAs across all electrodes and time points. This analysis revealed a significant main effect of Task, as well as a main effect of location. No significant main effect of orientation or interactions was observed. Since many other studies have examined the influence of location, we will focus on the main effect of Task.

The main effect of Task consisted of two significant spatiotemporal patterns that emerged more clearly in a *Grasp −Place* subtraction. Figure 6 depicts the temporal and spatial distribution of these 2 clusters, along with their surface topologies. The first cluster occurred during movement preparation and early initiation, extending from −390 to +150 ms relative to movement onset. This preparatory cluster involved a broad fronto-central to posterior electrode distribution, with a mean F value of 8.91 and the strongest effect over Poz at −200 ms, p < 0.001. The second cluster emerged after movement onset and extended from +120 to +1500 ms. This execution-related cluster also covered a broad fronto-central, parietal, and occipital electrode distribution, with a mean F value of 10.88 and the strongest effect at channel Cz at +520 ms, p < 0.001. Together, these results indicate that *Grasp* and *Place* movements elicited reliable ERP differences in time, during (a) late movement preparation and (b) subsequent movement execution.

**Figure 6:**
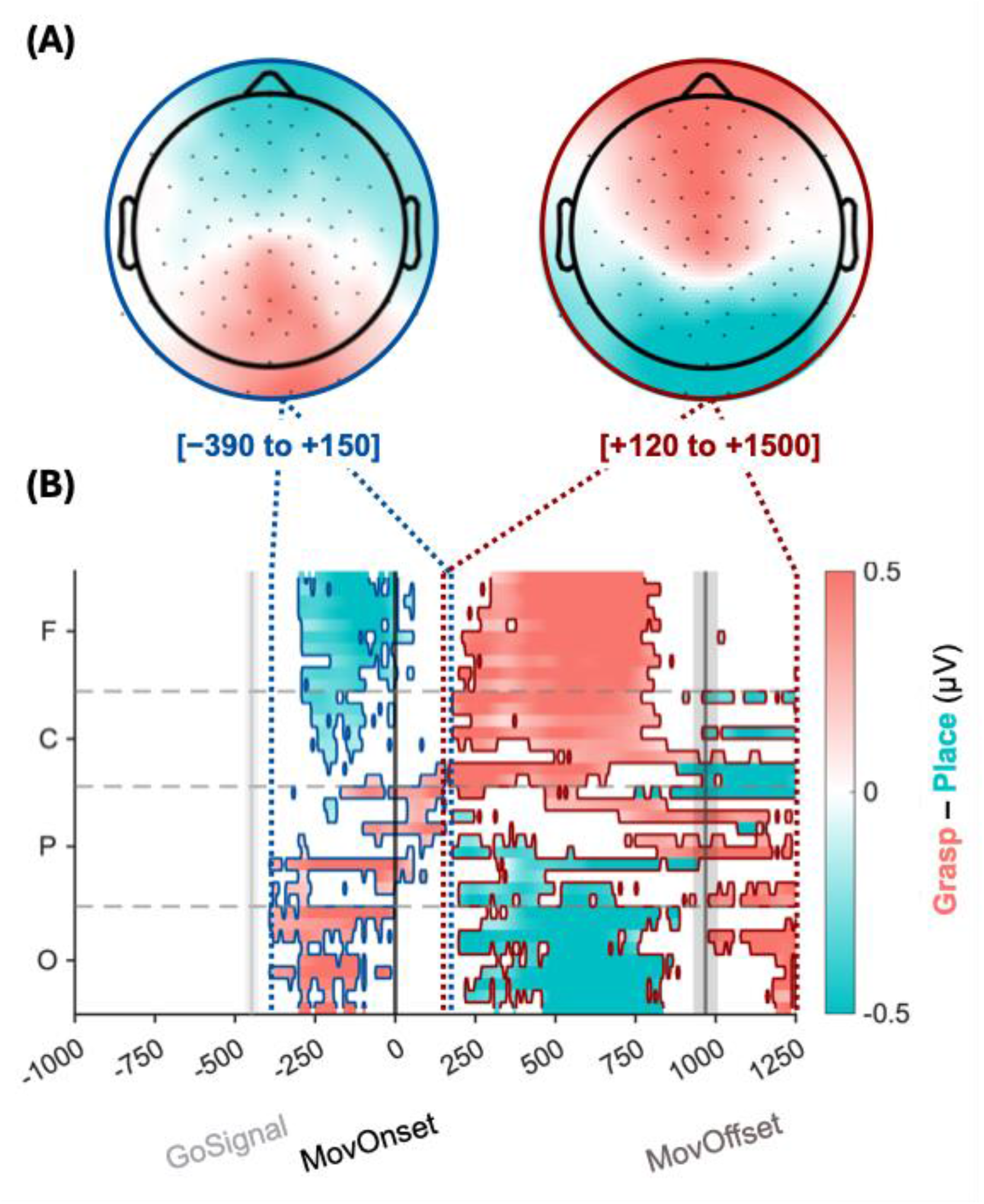
Mass-univariate LIMO analysis of the main effect of Task. **A**, Scalp topologies summarize the two significant spatiotemporal clusters: a preparatory cluster from −390 to +150 ms and an execution-related cluster from +120 to +1500 ms. The preparatory cluster (dark blue) showed relatively stronger fronto-central positivity and posterior negativity for *Place* compared with *Grasp*, whereas the execution-related cluster (dark red) showed the opposite polarity pattern, with a more sustained task difference after movement onset. **B,** The heat map shows the F-values for the task contrast between *Grasp* and *Place* across the same (F)rontal, (C)entral, (P)arietal, and (O)ccipital electrode families. Un-masked/colored regions indicate significant task-related differences identified by cluster-corrected repeated-measures ANOVA; the colourmap indicates the direction of the effect (teal for *Place*, pink for *Grasp*), and the contour colour indicates the cluster (dark blue for “Preparatory”, dark red for “Execution”).

Overall, the scalp distribution of the LIMO Task effect (Figure 6A) was consistent with the ‘switching’ pattern observed at movement onset in the ERP topologies of Figure 5, but showed that this pattern is stronger for the *Place* task: During the preparatory cluster, *Place* movements showed stronger fronto-central positivity and parietal-occipital negativity relative to *Grasp*. During the execution-related cluster, the polarity pattern reversed and was once again stronger but also more prolonged for *Place*, indicating a more sustained posterior positivity with corresponding fronto-central negativity. Thus, the Task effect primarily reflected changes in the strength and duration of shared ERP configurations rather than distinct scalp topologies.

CLUSTER AND SOURCE ANALYSIS: REACH MICROSTATES. To estimate the possible sources of our scalp ERPs, we first analyzed them using a Cartool microstate segmentation on movement-onset-locked ERPs from −500 to +500 ms (version 3.33; Brunet et al., 2011; http://brainmapping.unige.ch/Cartool.htm). This analysis identified five quasi-stable spatiotemporal scalp-level maps, present in both tasks, which together captured the topologic configurations that were dominant during the transition from movement preparation to execution (Figure 7A). Please note that the identified “microstate” maps are in close spatio-temporal agreement with the ones presented in the grand-average ERP topologies (Figure 5C), confirming that the hierarchical-clustering procedure successfully isolated quasi-stable representations that were already present in the data.

**Figure 7:**
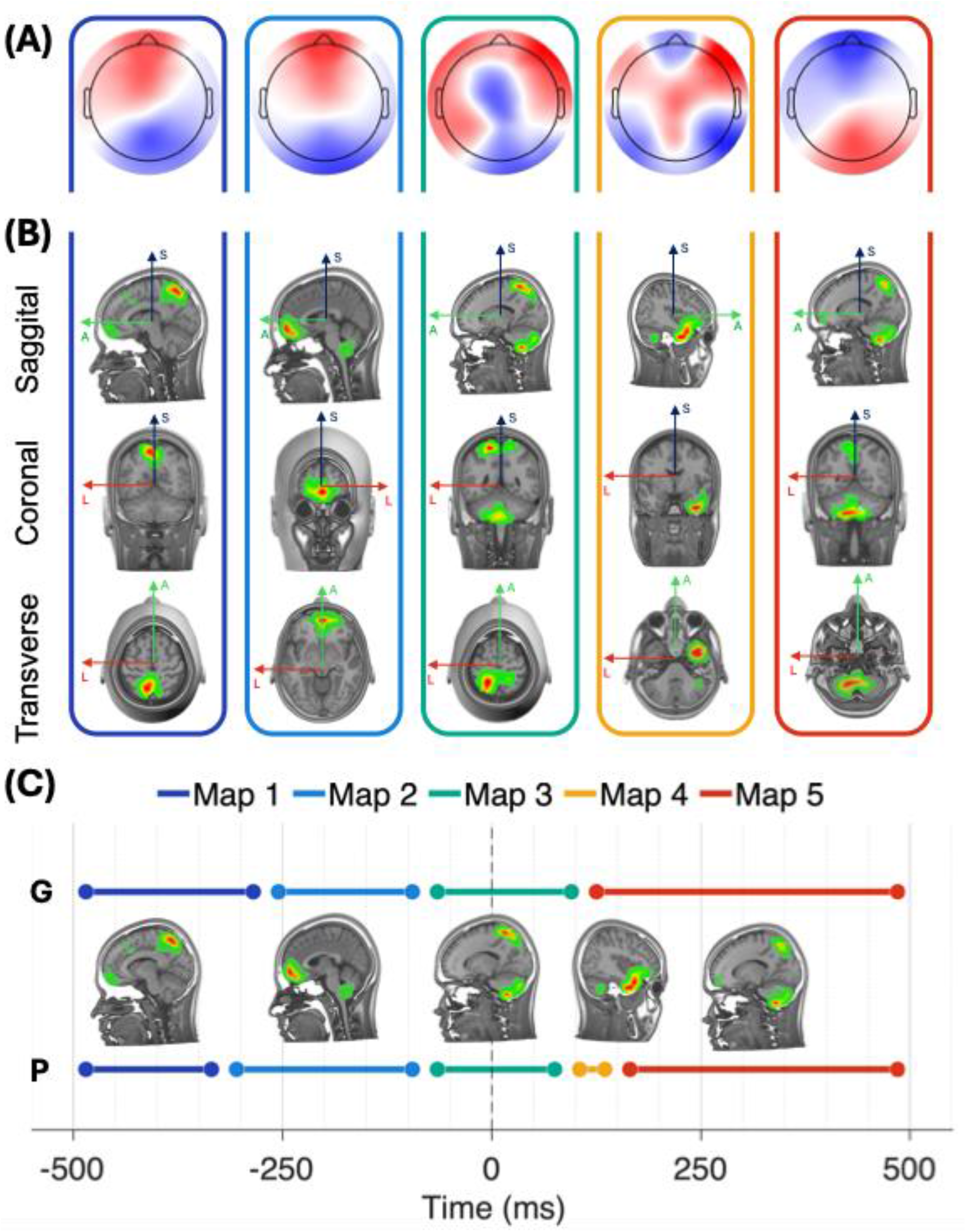
Spatial extent and temporal occurrence of the five microstate maps across the −500 to 500 ms movement-onset-locked epoch. **A**, Scalp topologies for each of the five template maps. **B**, Corresponding LAURA source estimates shown in sagittal, coronal, and transverse views of the distributed source estimate. Atlas labels and MNI coordinates indicate the three strongest regional maxima used for anatomical description. Map 1 was associated with precuneus, superior frontal cortex, and vmPFC sources; Map 2 with vmPFC, SMA, and cerebellum; Map 3 with precentral, postcentral, and cerebellar sources; Map 4 with ventral visual, lateral temporal, and precuneus sources; and Map 5 with cerebellum, precuneus, and vmPFC. Source estimates are shown descriptively to aid anatomical interpretation of the scalp-level microstate configurations. **C**, Coloured horizontal lines indicate the average temporal interval assigned to each map during back-fitting, with movement onset marked by the vertical dashed line, with *Grasp* (G) above and *Place* (P) below the source projections.

Next, we used LAURA to estimate the potential sources of these spatiotemporal scalp-level clusters. Figure 7B shows the resulting anatomical interpretations of the five template maps (Figure 7A; Table 1): Map 1 was associated with precuneus, superior frontal cortex, and ventral-medial prefrontal cortex (vmPFC) sources. Map 2 was associated with vmPFC, supplementary motor area (SMA), and cerebellar sources. Map 3 was associated with precentral (M1), postcentral (S1), and cerebellar sources. Map 4 was associated with ventral visual (Broadman area 20), lateral temporal, and precuneus sources. Finally, Map 5 was associated with cerebellar, precuneus, and vmPFC sources. Based on back-fitting to subject-level ERPs (Supplementary Figure 4), these maps occurred in partially overlapping temporal periods between the two tasks (Figure 7C): Map 1 from −426 to −123 ms, Map 2 from −34 to −1 ms, Map 3 from −88 to 32 ms, Map 4 from 72 to 156 ms, and Map 5 from −66 to 465 ms. Thus, Maps 1 and 2 primarily characterized preparatory activity, Map 3 spanned movement onset, and Maps 4 and 5 characterized early execution-related activity.

**Table 1.** AAL3 atlas-based source maxima for the five microstate template maps. For each microstate template map, the three strongest atlas-labelled source maxima are reported together with their corresponding Montreal Neurological Institute (MNI) coordinates. Source estimates were obtained using the AAL3 atlas and are presented descriptively to aid anatomical interpretation of the scalp template maps.

| Template map | Rank | Atlas region | MNI coordinates (x, y, z) |
| --- | --- | --- | --- |
| Map 1 | 1 | Precuneus | -9, -60, 60 |
| Map 1 | 2 | Superior frontal cortex | 28, 60, 9 |
| Map 1 | 3 | Ventromedial prefrontal cortex | 3, 60, -9 |
| Map 2 | 1 | Ventromedial prefrontal cortex | 3, 60, -9 |
| Map 2 | 2 | Supplementary motor area | 3, 22, 54 |
| Map 2 | 3 | Cerebellum | 22, -60, -54 |
| Map 3 | 1 | Precentral cortex | -35, -9, 60 |
| Map 3 | 2 | Postcentral cortex | 22, -41, 66 |
| Map 3 | 3 | Cerebellum | 22, -60, -54 |
| Map 4 | 1 | Ventral visual cortex | 28, 3, -41 |
| Map 4 | 2 | Lateral temporal cortex | 41, 3, -41 |
| Map 4 | 3 | Precuneus | -9, -72, 41 |
| Map 5 | 1 | Cerebellum | 22, -60, -54 |
| Map 5 | 2 | Precuneus | -9, -72, 41 |
| Map 5 | 3 | Ventromedial prefrontal cortex | 3, 60, -9 |

The presence of all five microstates in both *Grasp* and *Place* trials indicates that the two tasks engaged a shared sequence of large-scale neural configurations during the transition from movement preparation to execution. Thus, microstate segmentation supports the view that *Grasp* and *Place* rely on common neural mechanisms, identifies the potential neural substrates, and suggests that differences in these behaviors might arise from different timing and recruitment of the resulting segments.

MICROSTATE TASK SPECIFICITY. To confirm if *Grasp* and *Place* differed in the recruitment of our five fitted template maps, we analyzed the subject-level back-fitting measures extracted for each task and map. Specifically, repeated-measures ANOVAs were conducted on Time Coverage (%) and mean Global Field Power (GFP), with Task and Map as within-participant factors. Time coverage indexed the proportion of the peri-movement epoch during which each template map was dominant, whereas mean GFP indexed the strength of the scalp electric field during the periods assigned to that map.

Figure 8 shows task-specific differences in the duration and strength of scalp-level potentials for Map 2 during movement preparation and for Map 5 during movement execution. For the measure of Time-coverage (%) (i.e., the percentage of the input time-window, from −0.5 to +0.5 seconds relative to movement onset, that a certain map dominated), we observed a significant Task × Map interaction (F*(2.01,60.3) = 4.00,* p *= 0.023, ηp² = 0.118).* Follow-up paired comparisons showed that Map 2 occupied a greater proportion of the preparation epoch during *Place* than *Grasp* movements (Place: 29.2%; Grasp: 23.1%; *t*(30) = 2.54, FDR-corrected *p* = 0.041), whereas Map 5 occupied a greater proportion of the execution epoch during *Grasp* than *Place* movements (Grasp: 42.1%; Place: 39.3%; *t*(30) = −2.55, FDR-corrected *p* = 0.041). No other map showed a significant task difference in time coverage.

**Figure 8:**
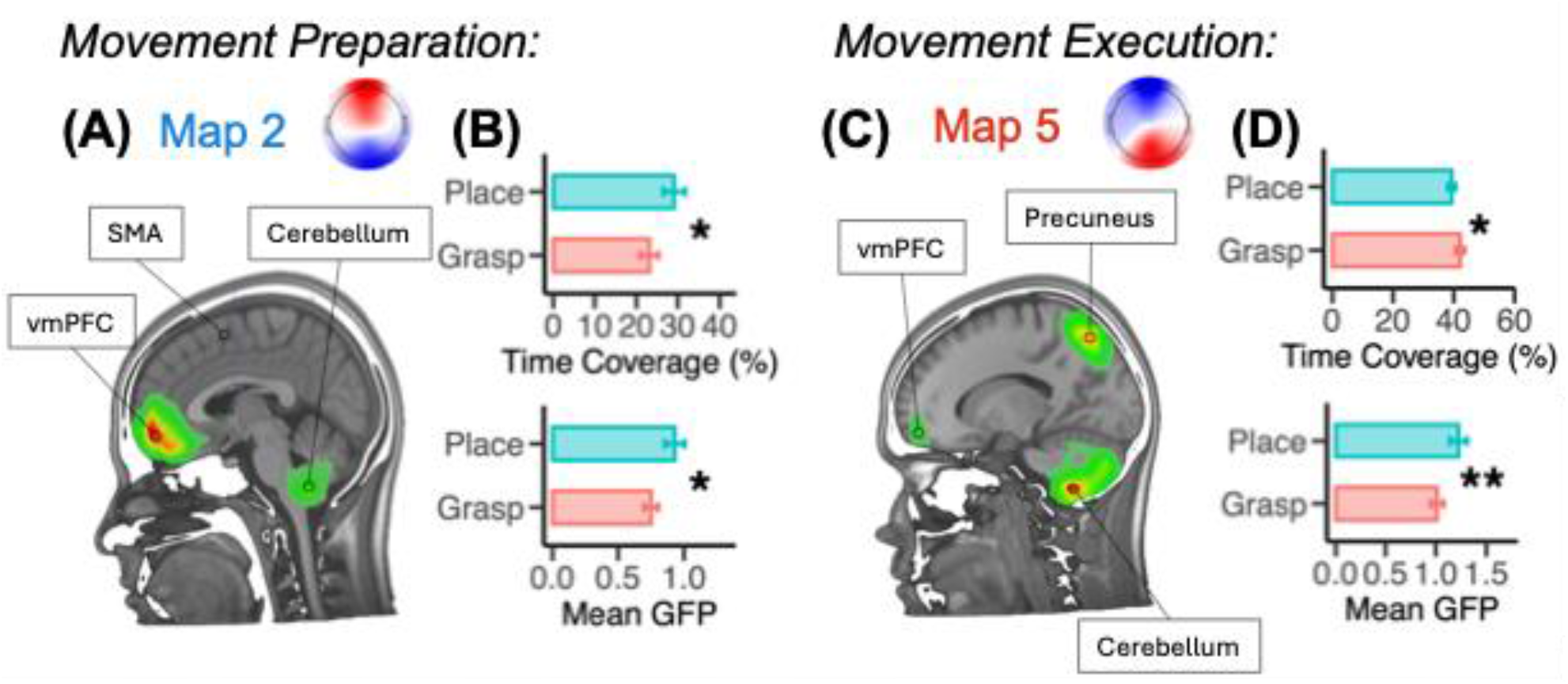
Task-sensitive microstate maps during movement preparation and execution. **A,** Distributed LAURA source estimates for the preparatory task-sensitive template Map 2, with the corresponding scalp topography shown in the upper right. Estimated sources were most prominent in vmPFC, SMA, and cerebellum. **B,** Task-specific back-fitting measures for Map 2. Bars show participant means, and error bars indicate SEM. Map 2 showed higher time coverage and mean GFP for *Place* than *Grasp*, indicating longer and stronger expression of this preparatory scalp configuration during placing. **C,** Distributed LAURA source estimates for the execution-related task-sensitive template Map 5, with the corresponding scalp topography shown in the upper right. Estimated sources were most prominent in the cerebellum, precuneus, and vmPFC. **D,** Task-specific back-fitting measures for Map 5. Map 5 showed higher time coverage for *Grasp* than *Place*, consistent with earlier and more sustained dominance in the *Grasp* ERPs, whereas mean GFP was higher for *Place*, indicating stronger scalp field magnitude during periods assigned to this map. Asterisks (*) indicate significant paired task comparisons after FDR correction.

Our measure of mean GFP (i.e., the strength of the electric field recorded at the scalp for the duration of a specific map) presented a main effect of Task (F*(1,30) = 16.57,* p *< 0.001, ηp² = 0.021)*, with *Place* producing overall stronger responses than *Grasp* across maps. Map-specific post hoc comparisons showed that mean GFP was significantly greater for *Place* than for *Grasp* for Map 2 and Map 5. For Map 2, *Place* showed higher GFP than *Grasp* (Place: 0.930; Grasp: 0.750), t(30) = −2.51, FDR-corrected p = 0.044, Cohen’s dz = −0.45. For Map 5, *Place* also showed higher GFP than *Grasp* (Place: 1.224; Grasp: 1.011), t(30) = −3.97, FDR-corrected p = 0.002, Cohen’s dz = −0.71. No significant task differences in mean GFP were observed for Maps 1, 3, or 4 after FDR correction. Overall, although only Map 2 showed greater temporal coverage for *Place*, both Map 2 and Map 5 exhibited stronger scalp field strength during *Place* movements.

Together, the microstate results confirm that *Grasp* and *Place* did not recruit entirely separate scalp configurations. Instead, the tasks differed in the temporal prevalence and strength of shared peri-movement maps, with *Place* showing increased recruitment of a preparatory vmPFC/SMA/cerebellar configuration and greater scalp field strength produced by cerebellar/precuneus/vmPFC regions during execution.

SPECTRAL ANALYSIS: TASK-SPECIFIC FREQUENCY SPECTRA. Next, we asked which frequency components carried the task-specific information identified by the preceding analyses. We examined frequencies from 1-30 Hz using overlapping 2-Hz passbands advanced in 1-Hz steps, without assuming that task information was restricted to the canonical frequency bands. We then used temporal-generalization analysis to test whether these task-discriminative patterns were time-specific, sustained, or re-expressed across different phases of movement preparation and execution.

Frequency-resolved SVM decoding first identified the spectral range carrying task-specific information (Figure 9B). A single significant cluster encompassed center frequencies from 2-6 Hz, corresponding to passbands spanning 1-7 Hz (cluster-corrected p < 0.001). Based on this low-frequency concentration, Figure 9A shows the corresponding time-resolved decoding profiles for the broadband signal and for low-(≤6 Hz) and higher-frequency (>7 Hz) components. The ≤6-Hz signal showed sustained above-chance decoding from −758 to +1993 ms (cluster-corrected p < 0.001), peaking at 74.55% at +602 ms. Broadband decoding showed a similar but weaker temporal profile, remaining significantly above chance from −558 to +1993 ms and peaking at 71.60% at +1112 ms. In contrast, the >7-Hz signal showed no cluster-corrected significant interval. The frequency × time analysis further localized when this low-frequency task information was expressed (Figure 9C). Together, these results show that the EEG information distinguishing *Grasp* from *Place* was predominantly expressed in slow-frequency activity, emerging during late movement preparation and persisting throughout execution.

**Figure 9:**
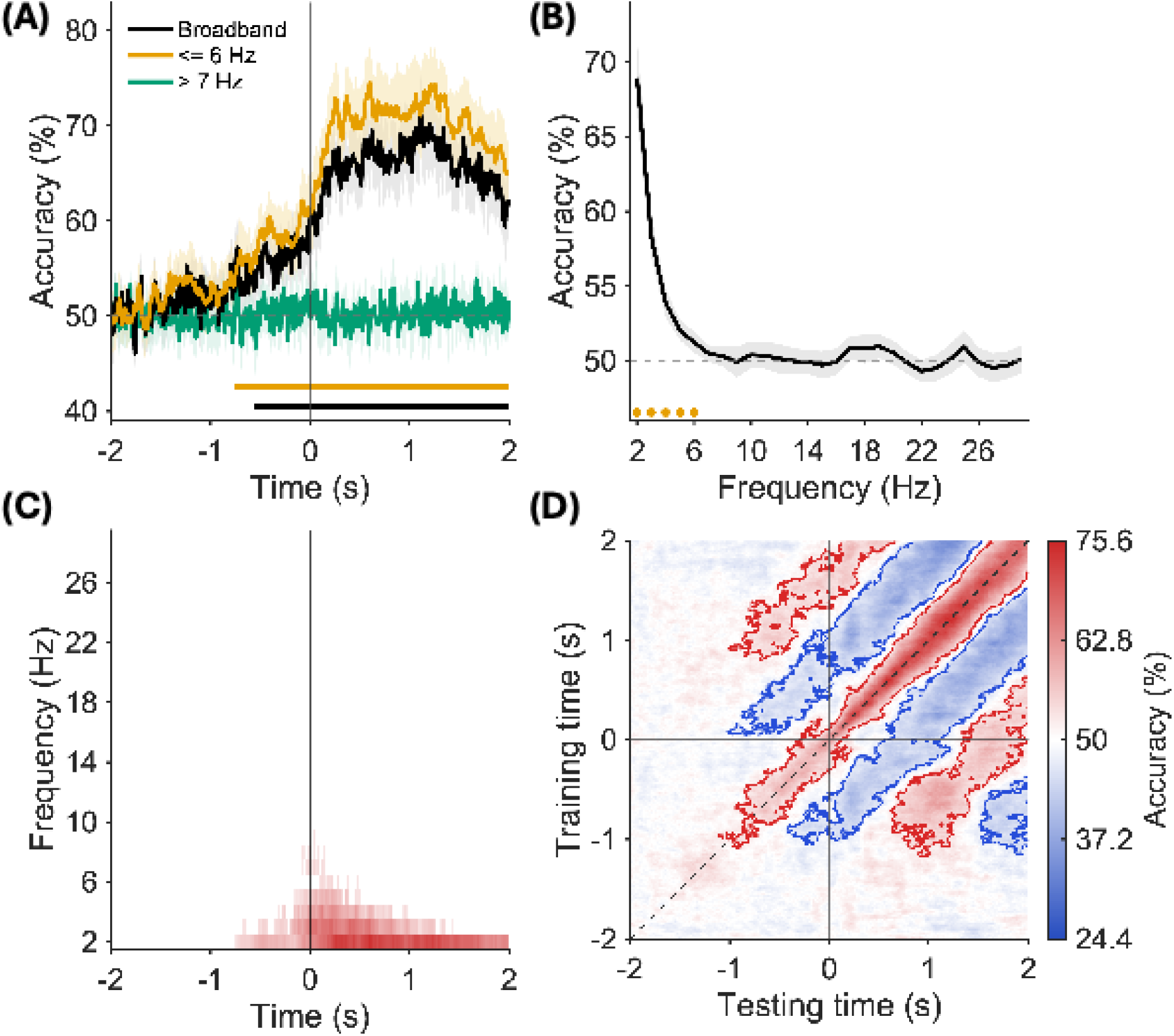
Frequency-specific and temporal-generalization decoding of *Grasp* versus *Place*. **A**, Time-resolved classification accuracy for broadband (black), ≤6-Hz (orange), and >7-Hz (green) EEG signals. Lines show group means and shaded regions indicate 95% confidence intervals. Horizontal bars indicate time intervals with cluster-corrected above-chance decoding for the corresponding signal; no significant interval was observed for >7 Hz. The vertical line marks the movement onset and the dotted horizontal line indicates chance performance (50%). **B**, Frequency-specific decoding accuracy for overlapping 2-Hz passbands advanced in 1-Hz steps. Shading indicates the 95% confidence interval, and orange markers indicate frequency ranges belonging to the significant (above-chance) cluster. **C**, Frequency × time representation of classification accuracy. Colored regions indicate frequency-time samples belonging to a significant cluster, whereas nonsignificant samples are displayed at chance (white mask). The vertical line marks movement onset. **D**, Temporal-generalization matrix for the ≤6-Hz signal. Classifiers were trained at each time point (y-axis) and tested at every other time point (x-axis). The dashed diagonal represents identical training and testing times, and vertical and horizontal lines indicate movement onset. Red and blue contours indicate significant above- and below-chance clusters, respectively, identified using two-tailed cluster-based sign-permutation tests.

EVOLUTION OF NEURAL REPRESENTATIONS. Finally, we employed a temporal generalization analysis to determine how the slow-oscillatory task-specific representation described in the last section evolved through time. Figure 9D shows the temporal-generalization matrix for the ≤6-Hz signal, illustrating how task-discriminative patterns generalized across movement preparation and execution. The dominant above-chance cluster was centered on the train-test diagonal and extended across training times from −1078 to +1993 ms and testing times from −1118 to +1993 ms (cluster-corrected p < 0.001). Two additional off-diagonal clusters linked temporally separated movement phases: classifiers trained from −1168 to +622 ms generalized to testing times from +632 to +1993 ms (p = 0.002), while classifiers trained from +802 to +1993 ms generalized back to −998 to +652 ms (p = 0.0095). Importantly, these above-chance regions alternated with significant below-chance off-diagonal bands, producing an oscillating or reversing temporal-generalization pattern (King and Dehaene, 2014), where the discriminative pattern changes polarity across time and causes classifiers to systematically reverse task labels at temporally displaced phases. Further details are provided in Supplementary Figure 5. Based on these data, we conclude that Task was predominantly represented by the slow-frequency oscillations (below 7Hz) spanning throughout movement preparation and execution, possibly reflecting the same parietofrontal polarity switch present in our ERP analysis (Figure 5).

## Discussion

Whereas our previous fMRI study compared the cortico-cerebellar topology of *Grasp* vs. *Place* (Luabeya et al., 2026) and found relatively subtle differences, the current study asks a novel and fundamental question: how are the spatiotemporal dynamics of such networks modulated to produce different behaviors? After confirming that our grasp and place tasks employed different spatiotemporal strategies (*Place* showing quicker reaction times, slower movements, and higher endpoint precision), we examined EEG data to identify the cortico-cerebellar substrates. At the level of ERPs, *Grasp* and *Place* shared a common peri-movement fronto-parietal polarity switch, but *Place* showed stronger, more prolonged preparatory and execution-related dynamics. Microstate and source analysis further traced this process to the differential recruitment of specific brain clusters through time. Finally, SVM decoding showed that task-specific information was carried mainly by an oscillatory low-frequency EEG component, evolving across movement preparation and execution. The following sections consider each of these topics in detail.

### Placing and Grasping involve different control strategies

Our first aim was to test if the *Place* and *Grasp* utilize different control strategies to meet their different computational demands. Specifically, feedforward control uses initial target information to specify a motor plan, whereas feedback control uses sensory information to update the ongoing movement command (Desmurget & Grafton, 2000; Shadmehr & Krakauer, 2008). These mechanisms are complementary, but weighted differently depending on task demands (Smeets & Brenner, 1999; Castiello, 2005; Fitts, 1954).

Compared to *Grasp*, *Place* movements were initiated sooner, unfolded more slowly, and produced lower endpoint grip-orientation variability (Figures 3, 4). The shorter reaction times in *Place* may be partly attributed to reduced feedforward demands (since the object was already held, participants did not need to estimate specific grip details), whereas the longer movement durations and lower peak velocities would optimize online visual feedback (for object alignment with the template).

### Place strengthens shared peri-movement ERP dynamics

Based on our previous study (Luabeya et al. 2026) and our behavioral data, we hypothesized that *Grasp* and *Place* use a similar mechanism that is dynamically modulated to produce these different movements. A prominent feature of the ERP results in both tasks was the reversal of the posterior-frontal polarity pattern around movement onset (Figure 5). Before movement onset, both tasks showed occipital-parietal negativity and fronto-central positivity, whereas after movement onset this pattern reversed. Similar EEG dynamics have been reported in studies, where preparatory activity before is followed by rapid changes in sensorimotor and parietal-occipital activity during execution (Tunik et al., 2008; Pereira et al., 2021; Xu et al., 2021). A comparable fronto-parietal switch was observed in our similar fMRI study (Luabeya et al. 20216), but here we observed the time course in real time, i.e., with a distinctive reversal exactly at movement onset. This reversal likely reflects a change in cortical states from prospective movement planning to execution-related sensory feedback and state estimation (Elsayed et al., 2016).

Although this pattern was present in both *Grasp* and *Place*, it was stronger during *Place*, likely reflecting the greater visual control demands imposed by the placement goal. During preparation, participants had to maintain not only the object’s location and orientation, but also object-template alignment. This may explain why *Place* amplified both preparatory (feedforward) activity and execution when sensory feedback occurs. Thus, our ERP data suggest increased strength and duration of an otherwise shared mechanism.

### Grasp and Place share a common microstate sequence

Our next aim was to identify quasi-stable brain states within our activity and trace the mechanism to specific brain regions. Our microstate analysis provided a topological description of how these scalp-level topologic states unfolded through time, and source localization provided anatomic estimates. Rather than identifying entirely separate task-specific maps, the segmentation again revealed a shared sequence of peri-movement states. Early preparatory maps involved regions associated with spatial goal representation, action planning, and prospective control, including precuneus, superior frontal cortex, medial frontal regions, SMA, and cerebellum, as expected for a network that transforms visual target information into action plans (Culham et al., 2003; Culham & Valyear, 2006; Blohm et al., 2009; Monaco et al., 2011; Crawford et al., 2011; Baltaretu et al., 2020).

The peri-onset involvement of precentral and postcentral sources is consistent with the transition from prospective planning to sensorimotor implementation, where motor commands and somatosensory consequences become more tightly coupled (Filimon et al., 2007; Begliomini et al., 2014; Fuehrer et al., 2026). Later cerebellar and parietal contributions are also consistent with predictive control and online updating during visually guided movement (Desmurget & Grafton, 2000; Miall & Wolpert, 1996; Shadmehr & Krakauer, 2008). Thus, the shared microstate sequence likely reflects a common visuomotor architecture that transitions from goal specification to movement initiation, to feedback-guided execution.

### Place modulates the timing and strength of shared microstates

Having established a common microstate sequence, we then asked how placement altered this sequence relative to grasping. Compared with *Grasp*, *Place* showed greater temporal coverage and stronger scalp field strength for the preparatory Map 2 (putative vmPFC/SMA and cerebellar sources), as well as stronger scalp field strength for the execution-related Map 5 (putative cerebellar, precuneus, and vmPFC sources). This pattern of results is consistent with changes in the weighting of a shared *Grasp*-*Place* architecture rather than recruitment of a separate placement network. Greater *Place* activation was estimated in Medial frontal cortex, implicated in voluntary action preparation, sequencing, and goal-dependent control (Cona & Semenza, 2017; Haggard, 2019; Wolpe et al., 2022), precuneus, associated with complex visuospatial transformations (Gorbet et al., 2004), and the cerebellum, implicated in visually guided online control and prediction of the sensory consequences of movement (Miall & Wolpert, 1996; Blakemore et al., 2001; Tseng et al., 2007). The cerebellum also showed greater activation for *Place* in our recent fMRI study (Luabeya et al., 2026).

### Low-frequency EEG dynamics distinguish Grasp from Place

Previous EEG/MEG studies have shown that reach planning and execution modulate activity across multiple frequency ranges (Van Der Werf et al., 2010; Blohm et al., 2019, 2022). Frequency-resolved multivariate decoding can reveal when and at which frequencies task-specific information is expressed (King & Dehaene, 2014; Bednar & Lalor, 2018). Here, our analysis showed that the *Grasp*-*Place* distinction was carried predominantly by low-frequency EEG activity, consistent with the previous results of movement-related EEG literature (Jahanshahi & Hallett, 2003; Shakeel et al., 2015), and with reach-to-grasp decoding studies (Sburlea et al., 2021; Pereira et al., 2021; Xu et al., 2021).

Finally, temporal generalization examined how low-frequency task-discriminative EEG patterns evolved across movement preparation and execution (King & Dehaene, 2014; Guo et al., 2019). The dominant diagonal cluster, surrounded by alternating above- and below-chance off-diagonal clusters, was consistent with an “oscillating or reversing” pattern (King & Dehaene, 2014). Below-chance generalization indicates that classifiers trained at one time point systematically predict the opposite task at another (Carlson et al., 2013), suggesting a reversal in the relative expression of task-discriminative features. This is consistent with the task-dependent modulation of the frontal-posterior polarity switch observed in our ERPs: compared with *Grasp*, *Place* showed stronger frontal positivity and posterior negativity during preparation, followed by stronger frontal negativity and posterior positivity during execution (Supplementary Figure 5). Thus, low-frequency task-discriminative features appear to reverse their relative expression across movement phases, rather than simply disappearing and reappearing.

### Limitations

Although EEG provides high temporal resolution, scalp recordings cannot localize neural generators with the precision of fMRI. Accordingly, our source estimates should be interpreted as estimates of regional constraints. It is reassuring that the putative regions localized based on our microstate analysis were generally aligned with our fMRI data (Luabeya et al. 2026), but may have underestimated subtle task differences. Second, EOG provided a useful EEG-compatible estimate of gaze-related timing but lacks the spatial precision necessary to entirely discount the influence of gaze on our EEG results. Third, the weighting of feedback vs feedforward movement control presented in our first hypothesis was inferred from behavioral data and task demands rather than direct experimental manipulation. Finally, the two actions differed in their initial hand state: *Grasp* began with an empty hand, whereas *Place* began with the object held. This is an inherent feature of grasping and placing, so we characterize our tasks as integrated sensorimotor behaviors, but it is difficult to dissociate the different contributions of cognitive, sensory, and motor signals to our results.

### Conclusions

This study investigated task-specific differences during planning and execution of *Grasp vs. Place* movements. *Place* altered movement timing and endpoint-orientation precision consistent with the increased visual demands of object-to-template alignment. These changes were reflected in our EEG data, where *Grasp* and *Place* shared a common parieto-frontal sequence of ERP and microstate configurations, but *Place* altered and strengthened preparatory and execution-related dynamics associated with cortical and cerebellar source estimates. Finally, decoding showed that task-specific information was best expressed in low-frequency EEG activity, below 6Hz, and unfolded dynamically across preparation and execution. While generally in spatial alignment with our previous fMRI study (Luabeya et al. 2026), these results reveal a key principle of visuomotor control: distinct motor goals can be supported by changes in the timing and strength of neural activity within a largely shared cortico-cerebellar network.

## Acknowledgments

The authors thank Dr. Xiaogang Yan and Saihong Sun for technical assistance. This research was funded by the Natural Sciences and Engineering Research Council of Canada Grant RGPIN-2022-04527. P.G. was supported by the Connected Minds program, funded by the Canada First Research Excellence Fund. The EEG equipment was acquired by P.J.K. using start-up funds from York University’s Vision: Science to Applications (VISTA) program, funded by the Canada First Research Excellence Fund. J.D.C. was supported by a York Research Chair.

## Author Contributions

P.G., G.N.L., E.F., P.J.K., and J.D.C. contributed to Conceptualization and Methodology. P.G. contributed to Investigation. P.G. contributed to Data curation, Software, Formal analysis, Validation, Visualization, and Writing −original draft. E.F., P.J.K., and J.D.C. contributed to Supervision. J.D.C. contributed to Resources, Project administration, and Funding acquisition. P.G., G.N.L., E.F., P.J.K., and J.D.C. contributed to Writing −review & editing.

## Data and Code Availability

Data and code are available upon request, subject to federal data-sharing policies, approval by the requester’s local ethics committee, and agreement on authorship for resulting publications.

## Supplementary Materials

### Power Analysis

Prior to completing data collection, a pilot-based power analysis was conducted using the datasets of the first 10 participants to determine the sample required to detect a *Grasp* −*Place* difference in the most prominent features in our data (Figure 5) and other recent EEG studies (Xu et al., 2021; Pereira et al., 2021). More specifically, the mean amplitude of theoretically justified ERP measurement windows and electrode regions (Keil et al., 2014), referring to the strength of the peri-movement frontal-posterior ERP reversal, was extracted as a dependent measure. For each participant and task, mean EEG amplitude was calculated over frontal (Fz, F1, F2) and posterior (Pz, P1, P2) electrodes during prespecified pre-movement (−300 to −100 ms) and post-movement (100 to 300 ms) intervals. A reversal score was then calculated as:

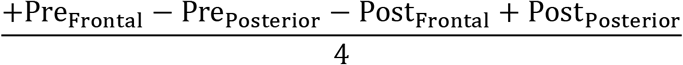

with the primary effect defined as the paired difference in this score between *Grasp* and *Place*. This one-degree-of-freedom contrast is equivalent to a Task (*Grasp, Place*) × Time (*Pre, Post*) × Region (*Frontal, Posterior*) interaction in a 2 × 2 × 2 repeated-measures design (Wiens & Nilsson, 2017). Using the pilot mean and standard deviation of the paired contrast, a two-sided paired-samples power analysis (α=.05, target power =.80) indicated that 30 analyzable participants were required (Supplementary Figure 1). Accordingly, the final sample of 30 participants met the prespecified recruitment target.

### Supplementary Figures

**Supplementary Figure 1:**
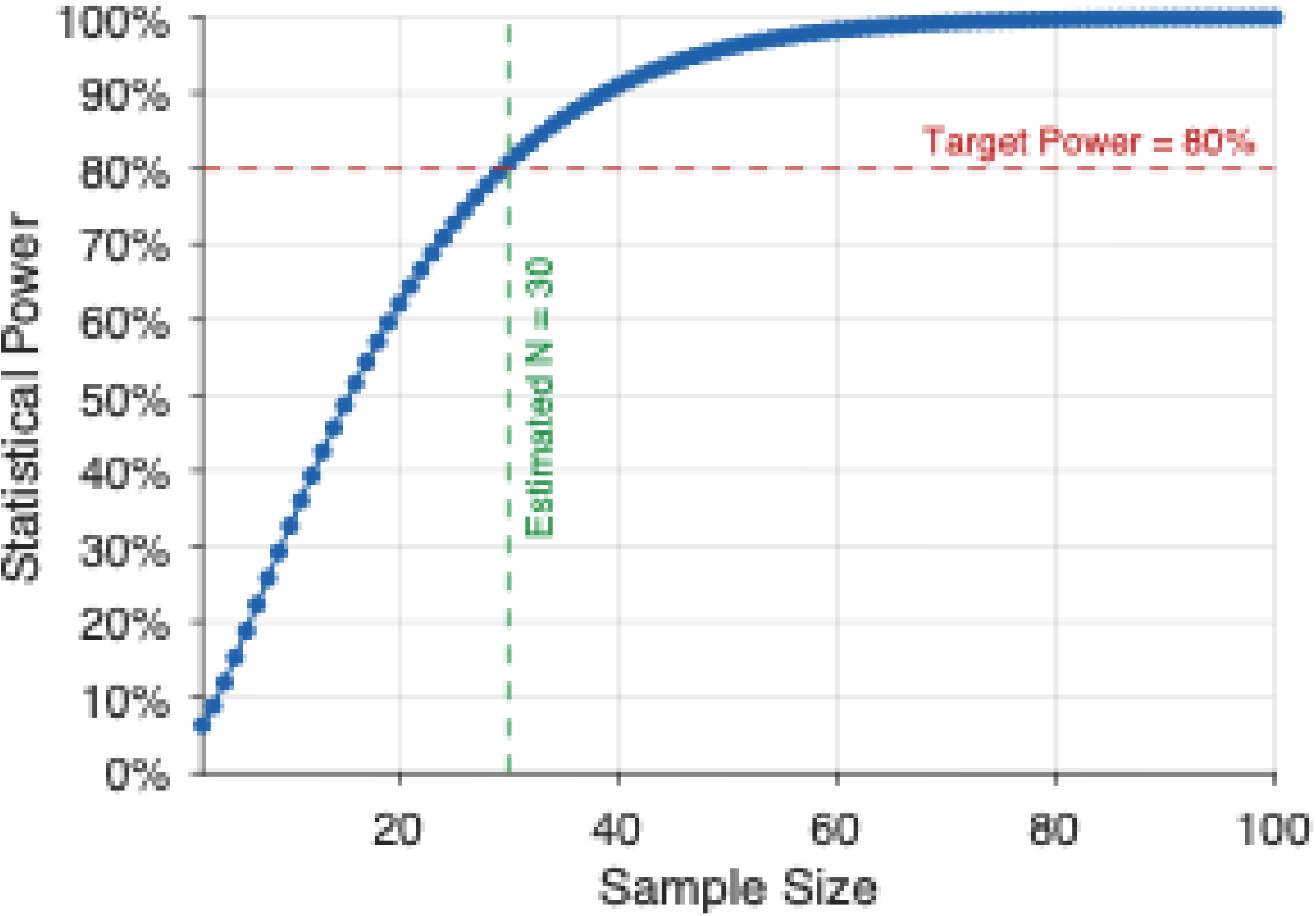
Pilot-based power analysis for the *Grasp*-*Place* difference in the peri-movement frontal-posterior ERP reversal. Statistical power for the one-degree-of-freedom Task × Time × Region interaction was calculated using the equivalent two-sided paired-samples contrast, with its mean and standard deviation estimated from the first 10 participants. The blue line and points show estimated power as a function of analyzable sample size. The horizontal red dashed line indicates the target power of 80%, and the vertical green dashed line indicates the minimum required sample of 30 participants (α=.05).

**Supplementary Figure 2:**
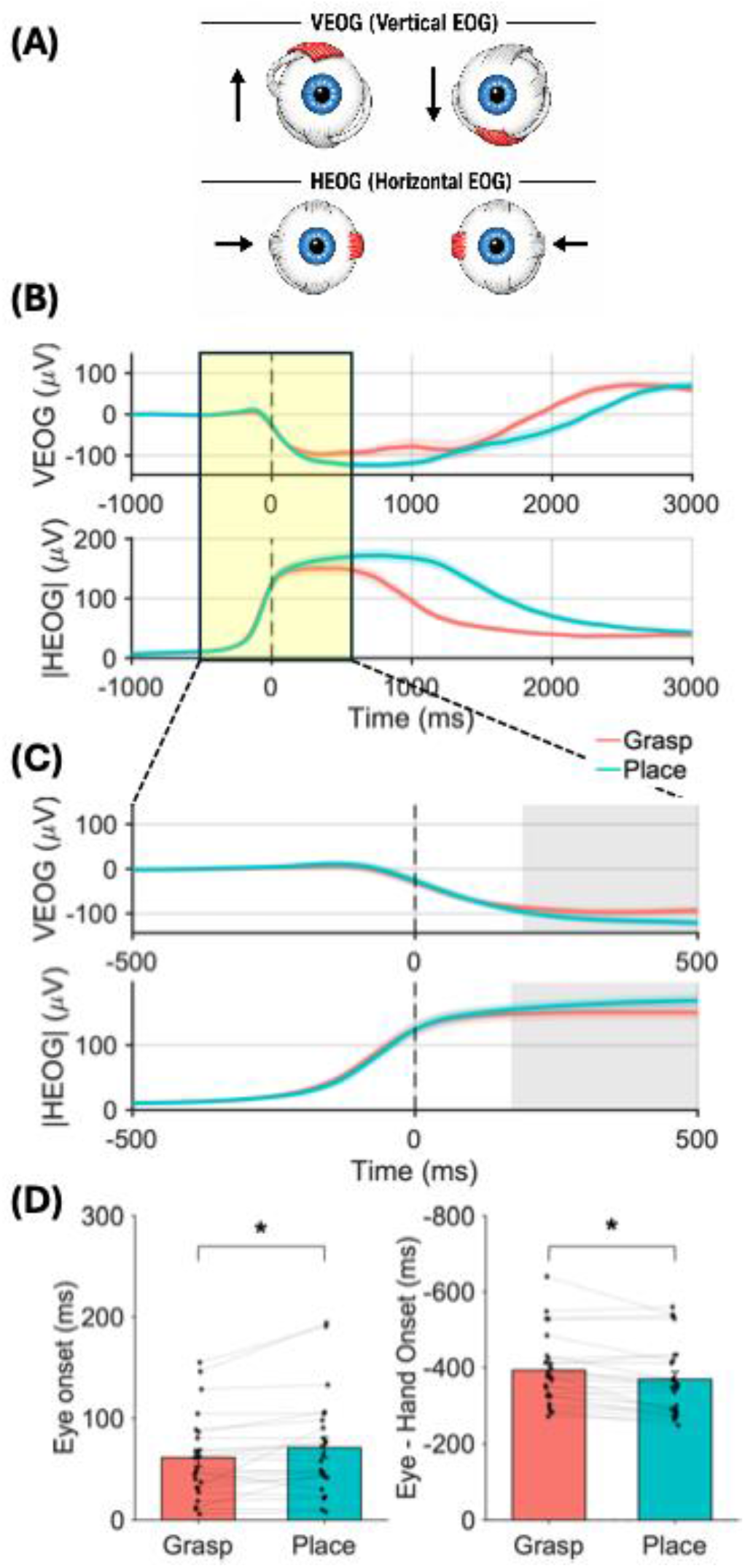
Eye-movement profiles during *Grasp* and *Place* movements. **A**, Schematic illustration of the vertical and horizontal eye-movement components estimated by the electrooculogram signals. Vertical EOG (VEOG) captures upward and downward eye movements, whereas horizontal EOG (HEOG) captures leftward and rightward eye movements. **B**, Grand-average movement-onset-locked VEOG and absolute HEOG (|HEOG|) traces for grasp and place trials, pooled across target location. Shaded regions around each trace indicate the SEM across participants. The vertical dashed line marks movement onset. The highlighted window indicates the interval expanded in C. **C**, Enlarged view of the peri-movement interval used for the task-effect analysis. Gray shaded regions indicate time windows in which grasp and place trials differed significantly after cluster-based correction. Red traces indicate grasp trials and cyan traces indicate place trials. **D**, Subject-level onset estimates for eye movements and their timing relative to hand movement onset. Left, mean eye-movement onset latency from the “GO”-signal for *Grasp* and *Place* trials. Right, mean eye-hand onset latency, computed as eye onset minus hand onset, with negative values indicating that eye movements preceded hand movement onset. Bars show the group means, error bars indicate SEM, black points show individual participants, and gray lines connect paired participant values across tasks. Asterisks indicate significant effects of task (p < 0.05).

**Supplementary Figure 3:**
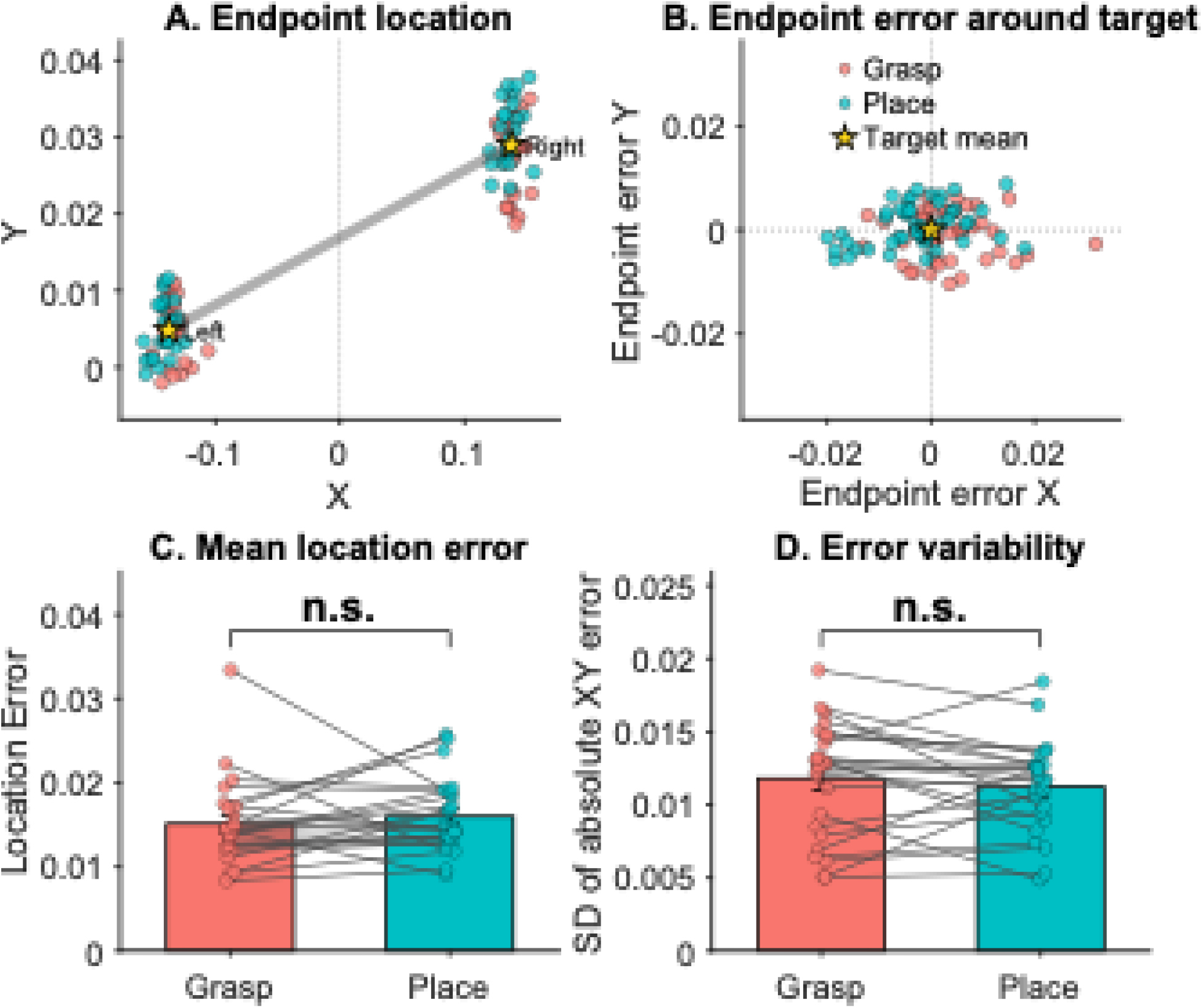
Endpoint-location error and variability for *Grasp* and *Place* trials. **A**, Subject-level endpoint locations for *Grasp* and *Place* trials, shown for the left and right target locations upon our experimental apparatus. Each point represents a participant × task × target-side mean endpoint location. Yellow stars indicate the estimated target (object/placeholder) location center. The vertical dashed line marks the centered midline. **B**, The same subject-level endpoint means expressed as target-centered error coordinates, with the target mean placed at the origin. **C**, Mean absolute error of the endpoint location for *Grasp* and *Place*, computed as the subject-level mean Euclidean error in the XY plane relative to the corresponding target center. **D**, Endpoint-error variability for *Grasp* and *Place*, computed as the subject-by-task standard deviation of the trial-level absolute XY endpoint error. In bar-graphs **C** and **D**, bars show group means, error bars indicate SEM, dots show individual participants, and gray lines connect paired observations from the same participant. *Grasp* trials are shown in red and *Place* trials in cyan. No significant task effects were observed.

**Supplementary Figure 4:**
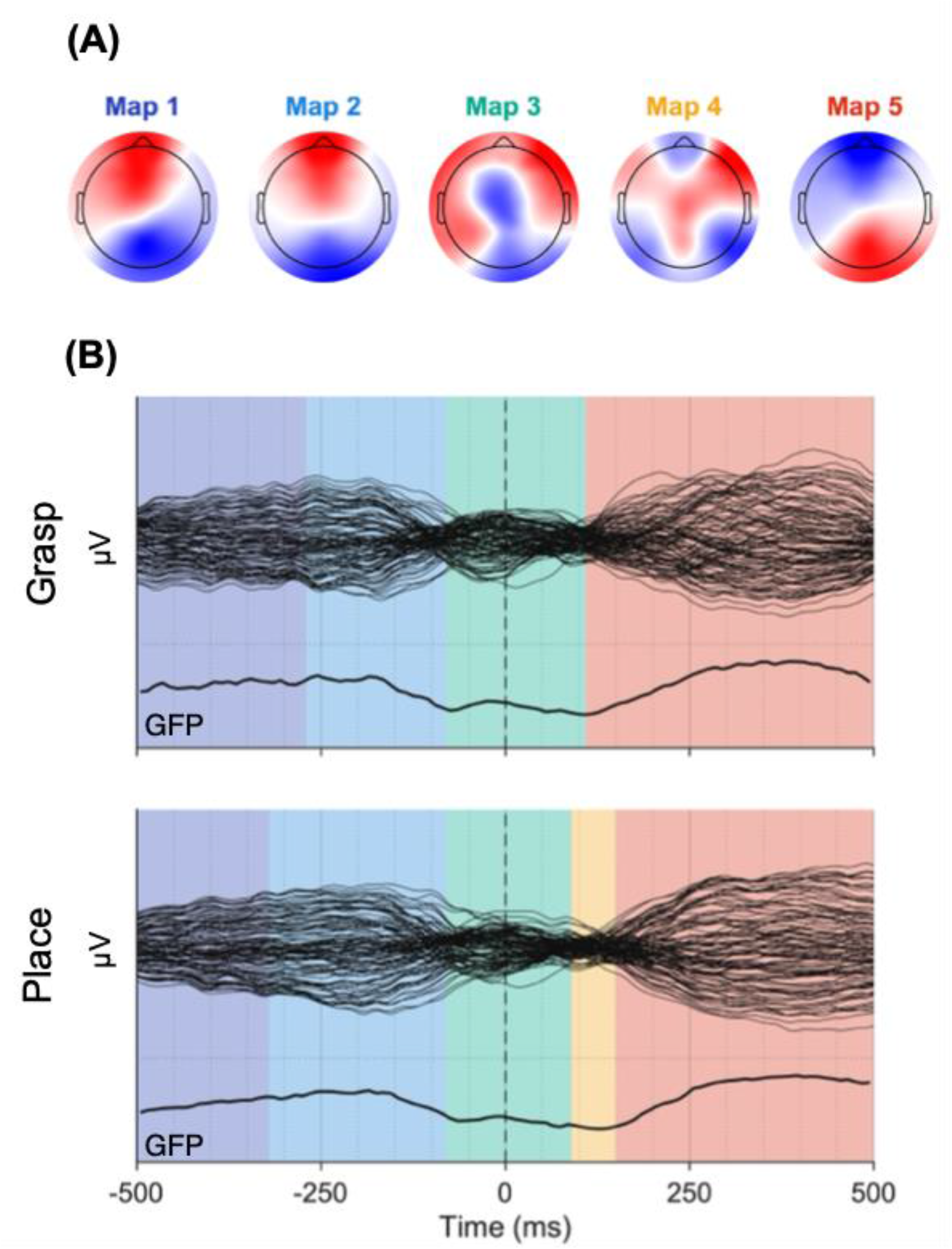
**A**, Scalp topologies of the five template maps identified by the microstate analysis. Colored map labels correspond to the temporal segments shown in panel B. Maps 1 and 2 primarily characterized preparatory activity before movement onset, Map 3 spanned the peri-onset transition, Map 4 appeared briefly after movement onset, and Map 5 dominated the execution-related period. **B**, Grand-average ERP topologies across the −500 to +500 ms peri-movement epoch for *Grasp* and *Place*. Black traces show electrode-wise ERP activity (butterfly plot), and the lower black trace shows the corresponding global field power (GFP). The vertical dashed line marks movement onset. Colored background segments indicate the temporal dominance of the five template maps (in A), fitted separately to the *Grasp* and *Place* ERPs. The segmentation shows that the two tasks expressed largely similar map sequences, but with differences in the timing and duration of specific maps.

**Supplementary Figure 5:**
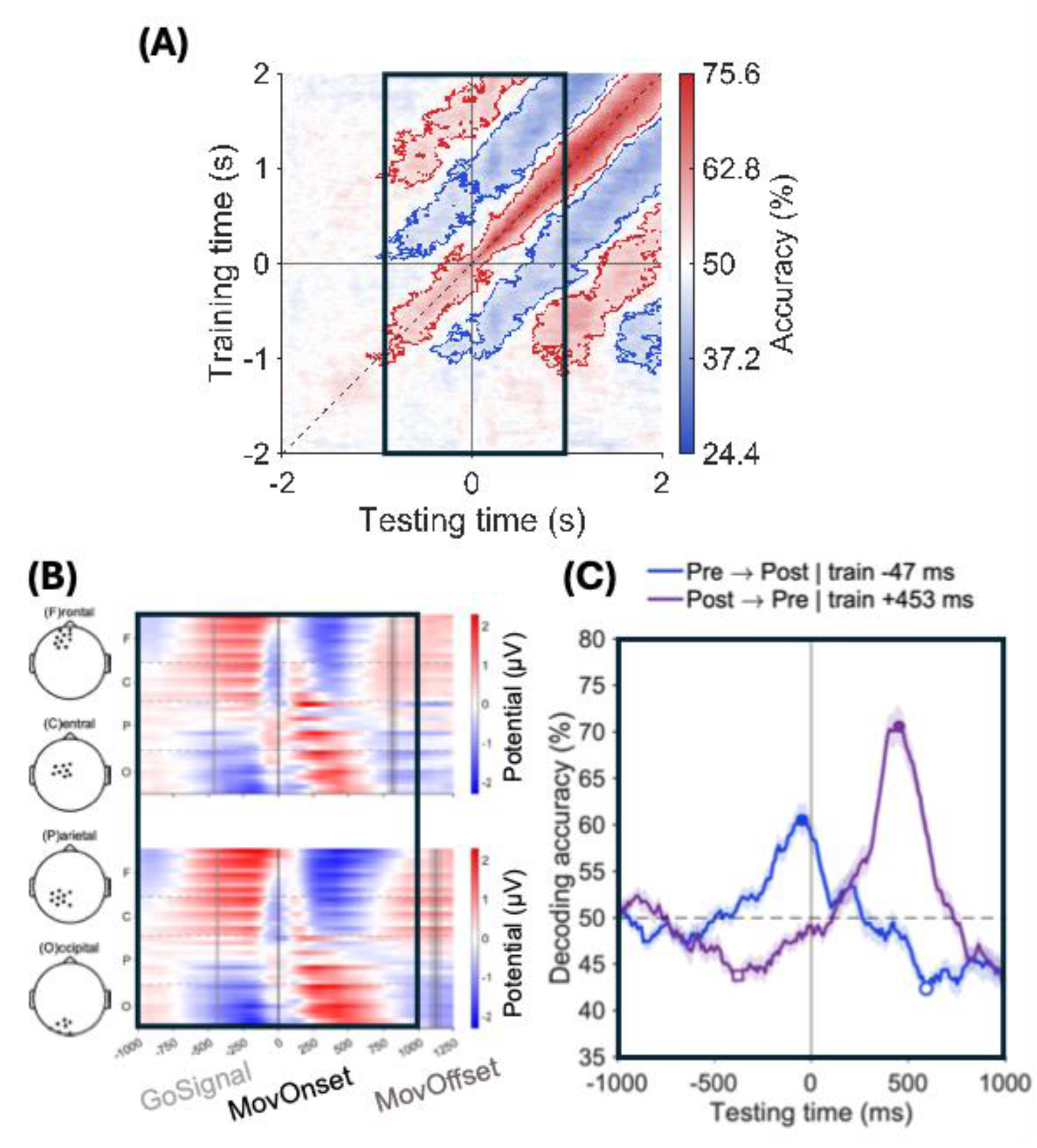
Low-frequency temporal generalization reveals a peri-movement reversal of task-discriminative EEG patterns. **A**, Temporal-generalization matrix for the ≤6-Hz EEG signal. Decoding accuracy is shown for classifiers trained at each time point (y-axis) and tested at every other time point (x-axis), relative to movement onset. Red contours indicate clusters with decoding significantly above chance, whereas blue contours indicate clusters significantly below chance, identified using two-tailed cluster-corrected sign-permutation tests across participants. The dashed diagonal denotes identical training and testing times, and the black box highlights the −1 to +1 s peri-movement interval examined further in B and C. The alternating above- and below-chance off-diagonal bands are consistent with an oscillating or reversing temporal-generalization pattern. **B**, Task-related frontal-posterior EEG potential patterns reproduced from Figure 5. During preparation, *Place* was characterized relative to *Grasp* by greater frontal positivity and posterior negativity, whereas this relative spatial pattern reversed during execution. This polarity reversal provides a plausible physiological basis for the below-chance cross-temporal decoding in A. **C**, Fixed-training-time decoding trajectories illustrating this peri-movement reversal. Classifiers were trained at the strongest significant negative cross-phase effects within the 01 to +1 s interval: −47 ms for the pre-to-post direction and +453 ms for the post-to-pre direction. Lines show mean decoding accuracy across participants, and shaded regions indicate SEM. Filled markers indicate decoding at the corresponding training time, whereas open markers indicate the selected cross-phase below-chance time point. The horizontal dashed line indicates chance performance (50%) and the vertical line denotes movement onset. Together, B and C show that the below-chance temporal-generalization pattern reflects a reversal in the relative expression of task-discriminative neural features across movement phases rather than a simple loss of task information.

